# Carbon: Decoding the Language of Life

**DOI:** 10.64898/2026.05.22.727119

**Authors:** Loubna Ben Allal, Qiuyi Li, Maurizio Fiusco, Lewis Tunstall, Kashif Rasul, Ed Beeching, Dana Aubakirova, Carlos Patiño, Thibaud Frere, Anton Lozhkov, Georgia Channing, Thomas Wolf, Diego di Bernardo, Leandro von Werra

**Affiliations:** Hugging Face; Zhongguancun Academy; TIGEM; University of Naples Federico II

## Abstract

Genomic foundation models have emerged alongside the rapid progress of large language models, offering a promising framework for learning general-purpose sequence priors for DNA understanding, generation, and design. This connection to LLMs creates a major opportunity: modern architectures, scaling infrastructure, autoregressive training, and token-based modeling provide powerful tools for genomic sequence modeling. At the same time, DNA differs fundamentally from natural language. Genomic sequences are noisy, redundant, sparsely constrained, unevenly annotated, and shaped by evolutionary rather than communicative pressures. As a result, key components of the standard LLM recipe, including data construction, tokenization, and training objectives, must be reconsidered in the biological sequence setting. A central challenge in DNA modeling is reconciling single-nucleotide resolution with long-context reasoning. Single-nucleotide resolution is essential for variant effect prediction, splice-site analysis, and codon-level reasoning. Long-context modeling is equally important, as many genomic mechanisms depend on distal regulatory elements, gene neighborhoods, and long-range evolutionary constraints. However, the most direct path to nucleotide-level reasoning, single-nucleotide tokenization, makes genomic sequences extremely long and imposes substantial computational cost on Transformer models. We present Carbon, a family of efficient generative DNA language models designed as a practical reference point for this setting. Carbon includes 3B- and 8B-parameter decoder-only autoregressive models using non-overlapping 6-mer tokenization. Carbon-3B supports a maximum context length of 65,536 tokens, corresponding to approximately 393 kbp of DNA; Carbon-8B supports up to 131,072 tokens, roughly 786 kbp. This simple and controlled setup helps isolate a central question for DNA language modeling: whether current progress is limited primarily by model architecture and nominal context length, or by more basic alignment between data, tokenization, objectives, evaluation, and the biological structure of genomic sequence. In our training-free evaluation suite, Carbon-3B is competitive with Evo2-7B despite having less than half the parameters. Carbon-8B improves on Carbon-3B on every training-free task, with the largest gain on long-context retrieval. Both models deliver tens-fold faster inference under comparable settings. The Carbon recipe combines annotation-aware data curation, deterministic 6-mer tokenization, and a staged CE-to-FNS objective schedule, adapting the LLM recipe to the statistical and biological properties of DNA rather than directly transplanting it. We release the models, data, training code, and evaluation suite, including new training-free probes for sequence-level perturbation and DNA long-context retrieval. Carbon is intended as an open recipe for efficient generative DNA modeling rather than an argument for any specific architecture, tokenization strategy, or objective design as the optimal solution. Its strong performance provides grounded evidence that substantial room remains for domain-aware model design carefully aligned with the genomic sequence itself.

## 1 Introduction

Genomic DNA encodes protein-coding genes, regulatory elements, non-coding RNAs, repetitive structures, and evolutionary constraints accumulated over long evolutionary timescales [1]. A foundation model that can learn the distribution of genomic sequence has the potential to support a broad range of biological applications, including sequence completion, variant interpretation, functional annotation, comparative genomics, and programmable DNA design. Inspired by the rapid progress of large language models [2, 3, 4] and protein language models [5, 6], genomic foundation models have become an increasingly active direction for learning general-purpose sequence priors directly from DNA [7, 8, 9, 10, 11].

In this technical report, we present Carbon, a family of efficient generative DNA language models. Carbon includes 3B- and 8B-parameter decoder-only autoregressive models with non-overlapping 6-mer tokenization. The architecture follows a standard open-weight Transformer design (RMSNorm, SwiGLU, RoPE, GQA), as described in Section 5.1. Carbon-3B supports a maximum context length of 65,536 tokens (approximately 393 kbp of DNA), and Carbon-8B extends to 131,072 tokens (approximately 786 kbp). While recent genomic models often use long-convolution, state-space, or hybrid sequence architectures [12, 13, 14, 9], Carbon shows that a standard Transformer backbone combined with compact 6-mer tokenization can achieve strong performance while supporting long-context inference with substantially higher throughput. This choice also makes Carbon easier to use with familiar opensource LLM tools, and keeps the empirical focus on the components we ablate directly: data curation, tokenization, and the training objective. Across our training-free benchmarks, Carbon-3B is competitive with Evo2-7B [9] at less than half the parameter count. Carbon-8B beats Evo2-7B on the majority of variant-effect prediction, sequence generation, and perturbation tasks, and leads on long-context retrieval at 786 kbp. Both models run tens-fold faster at inference.

To build Carbon, we studied how the standard LLM recipe should be adapted for DNA. The most important differences we found arise at three levels: data, tokenization, and the training objective. The first difference is the signal-to-noise structure of genomic sequences. Unlike natural language text, which is produced for communication and is therefore relatively compact and information-dense, genomes are the product of evolutionary processes shaped by mutation, genetic drift, selection, duplication, transposition, and evolutionary contingency. Large portions of raw genomic sequence can be weakly constrained, repetitive, or only sparsely informative for the biological capabilities we want the model to acquire. This motivates annotation-aware data curation [10, 9] rather than treating raw nucleotide count as the only measure of data quality.

The second difference is the absence of stable word boundaries. BPE and related subword tokenizers [15] are powerful in natural language because repeated character patterns often correspond to reusable lexical or sublexical units. DNA contains motifs, but these motifs are often degenerate, context-dependent, strand-sensitive, length-variable, and tolerant to substitutions or insertions [16]. In our experiments, this made learned BPE tokenization poorly matched to autoregressive DNA generation. Non-overlapping 6-mer tokenization [10, 17] provided a more stable alternative: it fixes the prediction step length, avoids segmentation ambiguity, reduces token length by a factor of six (compared to a single-nucleotide tokenizer), and substantially improves inference efficiency with little performance cost.

The third difference is the structure of the training objective. Standard next-token cross-entropy works remarkably well in natural language modeling, where tokenizer units often behave as meaningful prediction targets. In a 6-mer DNA model, however, cross-entropy treats each 6-mer as an atomic class among 4,096 possibilities. This creates an all-or-nothing supervision signal: a prediction that matches five out of six nucleotides is penalized in the same way as a completely unrelated 6-mer. During Carbon training, we found that this exact-token objective can lead to a persistent “loss staircase” phenomenon and increased sensitivity to numerical precision in late-stage training. Factorized Nucleotide Supervision (FNS, [17]) addresses this problem by exposing nucleotide-level supervision through probability marginalization, providing structured partial credit while preserving the efficiency of 6-mer tokenization. In practice, switching from cross-entropy to FNS at the appropriate stage stabilized training and allowed the model to continue improving.

Together, these observations suggest that DNA language modeling is not simply natural language modeling with a four-letter alphabet. The standard LLM recipe remains extremely valuable as a starting point, including the use of largescale data, token-based sequence representation, and next-token cross-entropy. However, these components need to be reconsidered in the biological sequence setting. For Carbon, this led to an annotation-aware mixed corpus, focused primarily on eukaryotic genes, non-overlapping 6-mer tokenization, and a staged CE-to-FNS objective schedule.

Developing this recipe required ablations guided by evaluations that are stable, inexpensive, and directly comparable across pre-training runs. DNA foundation models are commonly evaluated through a mixture of zero-shot likelihood-based benchmarks, supervised finetuning tasks, and embedding-based analyses [9, 10, 11]. Each evaluation mode emphasizes a different aspect of model behavior: finetuning benchmarks measure task-specific transfer, embedding analyses probe learned representations, and zero-shot likelihood-based tasks test the base model directly without additional training. For pre-training ablations and base-model comparison, we focus on zero-shot, training-free evaluations, since they avoid confounding factors introduced by finetuning protocols or embedding classifiers. To select reliable evaluation tasks, we follow criteria used in recent LLM evaluation work [18]: a useful task should score above random, improve with training, exhibit low seed-to-seed noise, and produce broadly consistent model rankings across checkpoints. The resulting suite spans generation (Sequence Recovery [10]) and variant-effect prediction (ClinVar coding and non-coding [19, 20], BRCA2 [21], TraitGym [22]). To cover capabilities not addressed by existing public benchmarks, we additionally introduce two sequence-level perturbation probes (nucleotide triplet-expansion and synonymous codon replacement) and Genomic-NIAH, a DNA long-context retrieval benchmark built from real genomic haystacks.

The performance of Carbon suggests that the field is still far from a settled optimum: a relatively standard Transformer can match much larger and longer-context baselines. This means that there is substantial room for progress through careful algorithmic and data-centric work, not only by increasing the parameter count or the nominal context length. Several questions remain open, including how to construct genomic pre-training data; how to represent DNA sequences; how to recover nucleotide-level reasoning efficiently; how to measure effective long-context utilization; and how to connect sequence-level models to practical biological applications.

More broadly, Carbon reinforces the view that DNA language modeling should not be treated as a direct transplant of natural language modeling. LLM technology provides powerful tools, but DNA sequence is governed by different statistical and biological principles. The purpose of this report is therefore not only to introduce a strong model family, but also to share the implementation path, empirical observations, and design considerations behind it. We hope Carbon can serve as a grounded baseline for efficient generative DNA modeling and help clarify where the field currently stands, what a strong practical baseline can look like, and which questions may matter most for future progress.

### Main contributions

The main contribution of this report is an open, evaluation-driven recipe for efficient generative DNA modeling. We release the resulting models, training data, training code, and evaluation suite, together with ablations showing how data, tokenization, and the training objective shape performance.

- **Strong open genomic models**. We release Carbon-3B (393 kbp context) and Carbon-8B (786 kbp context). Carbon-3B is competitive with Evo2-7B [9] despite less than half the parameter count; Carbon-8B improves on Carbon-3B on every training-free task, with the largest gain on long-context retrieval. Both run tens-fold faster at inference.
- **Ablation-driven data recipe**. We construct an annotation-aware public-data mixture centered on annotated eukaryotic functional regions and ablate the key choices behind it: corpus source, data processing, metadata conditioning, and data mixing ratios.
- **Tokenizer and training objective for DNA**. We compare learned BPE and fixed 6-mer tokenization, and introduce a staged loss switch from cross-entropy to Factorized Nucleotide Supervision [17], making compact 6-mer modeling more stable and effective.
- **Open training-free evaluation suite with new DNA probes**. We release an evaluation suite spanning sequence generation, variant-effect prediction, controlled perturbation, and long-context retrieval. The suite introduces two new sequence-level perturbation probes, nucleotide triplet-expansion and synonymous codon replacement, and Genomic-NIAH, a DNA long-context retrieval benchmark built from real genomic haystacks, with tasks of varying difficulty up to 786 kbp.

## 2 Related Work

The development of DNA foundation models has followed several complementary paths. Early self-supervised models were largely encoder-only masked language models, including the DNABERT series [7, 23], the Nucleotide Transformer series [8], and the GENERanno series [24], which learned contextual sequence representations and achieved strong performance on supervised downstream prediction tasks. More recently, generative DNA models have become increasingly important, including autoregressive models such as megaDNA [25], HyenaDNA [12], Evo series [26, 27, 9], and the GENERator series [10, 17], as well as diffusion-based approaches such as MDLM [28] and DNA-Diffusion [29]. In parallel, supervised sequence-to-function models such as Enformer [30], Borzoi [31], and AlphaGenome [14] have demonstrated the value of directly predicting regulatory tracks, gene expression, splicing, and other functional genomic outputs from DNA sequence. Together, these models have established DNA foundation modeling as a rapidly evolving field at the intersection of representation learning, generative modeling, and functional genomics.

Across these developments, two goals have repeatedly emerged as central: single-nucleotide resolution and long-context modeling. Single-nucleotide resolution is essential because many biological questions depend on individual bases, including SNP interpretation, variant effect prediction, splice-site analysis, codon-level reasoning, and regulatory mutation scoring. Long-context modeling is equally important because many genomic mechanisms are nonlocal: distal regulatory elements can influence gene expression across large genomic distances, gene neighborhoods can carry functional organization, chromatin structure can couple distant loci, and evolutionary constraints may only become visible with sufficient surrounding context. An ideal DNA foundation model would therefore reason at the level of individual nucleotides while also integrating information over long genomic spans.

These two goals, however, are in tension. The most direct way to preserve single-nucleotide resolution is singlenucleotide tokenization, where each base is represented as one token. This choice makes nucleotide-level prediction natural, but it also makes encoded DNA sequences extremely long. For a Transformer model, the attention cost grows quadratically with token length, so increasing context length under single-nucleotide tokenization quickly becomes computationally expensive. This resolution–context trade-off is one of the core difficulties of DNA language modeling.

One line of work addresses this trade-off by preserving single-nucleotide tokenization while adopting more efficient architectures [32, 33]. This includes state-space and related long-sequence models such as Caduceus [13] and HybriDNA [34], Hyena-style architectures such as HyenaDNA and the Evo series, and encoder or sequence-to-function models that combine attention with convolutional or U-Net-like components [35], including Enformer, Borzoi, AlphaGenome, and NT-v3 [36]. These approaches are valuable because they retain base-level input resolution and reduce the cost of long-sequence processing. At the same time, efficient long-sequence architectures do not automatically guarantee effective long-context utilization. A model may support a long nominal input window, but the practical value of that context depends on whether additional sequence is converted into measurable improvements in prediction, generation, or downstream biological performance. In generative settings, inference cost is also important because likelihood scoring and autoregressive generation must often be repeated over many sequences, variants, or perturbations.

A second line of work keeps the Transformer architecture closer to standard language models but compresses the input sequence through coarser tokenization. Examples include fixed-length *k*-mer tokenizers [8, 10], learned BPE tokenizers [23, 37, 38], and other learnable compression schemes such as vector-quantized tokenization [39, 40]. This route is computationally attractive: grouping multiple nucleotides into one token reduces sequence length, improves context coverage under a fixed token budget, and can substantially reduce attention and decoding cost. However, coarse tokenization appears to create a different problem. If the model predicts one *k*-mer or learned token at a time, single-nucleotide probabilities are no longer directly available. This is a serious limitation for genomic applications, because nucleotide-level reasoning is not a peripheral requirement but one of the central use cases of DNA foundation models.

GENERator-v2 provided an important perspective on this problem: single-nucleotide tokenization is not the only way to obtain single-nucleotide resolution [17]. A coarse *k*-mer token is not a biological word; it is a computational compression unit with explicit internal nucleotide structure. If a model predicts a distribution over *k*-mer tokens, nucleotide-level probabilities can be recovered by marginalizing over all tokens that share the same base at a given position. Factorized Nucleotide Supervision formalizes this idea by exposing nucleotide-level learning signals from coarse-token logits. Under a conditional-independence approximation, a 6-mer model can be viewed as predicting six nucleotide distributions from a shared hidden state, while retaining the efficiency benefits of compressed tokenization. This creates a possible bridge between coarse tokenization and single-nucleotide reasoning.

This idea opened a useful direction, but it also left important questions unresolved. The original evidence was not yet sufficient to determine whether coarse-token generative modeling could scale into a strong alternative to singlenucleotide long-context models. The model scale was still limited, the ablations did not fully isolate the effects of data, tokenizer, and objective design, training stability was not extensively analyzed, and comparisons against much larger modern baselines remained incomplete. In other words, GENERator-v2 showed that the door was open, but it did not yet establish how far this route could go. Carbon is designed to provide a more grounded answer to this question by scaling the model, strengthening the evaluation, and systematically analyzing the components of the coarse-token generative recipe.

## 3 Data

Eukaryotic genomes are long sequences of chemical letters (bases) containing relatively sparse functional elements. In humans, the genome spans roughly 3 × 10^9^ base pairs, yet only about 1–2% directly encodes proteins, the molecular machines that perform most cellular functions [41]. Beyond protein-coding exons, a larger but still limited fraction shows evolutionary constraint, often reflecting regulatory, transcript-associated, or other non-coding functions [42]. Roughly 50% of the genome consists of repetitive sequences, many derived from ancient transposable elements that copied themselves throughout the genome over evolutionary time [41]. Much of the remaining sequence has no clearly established function. This creates a central challenge for genomic language modeling: biological signal is sparse, heterogeneous, and often difficult to separate from background sequence. We describe this as a low signal-to-noise ratio of raw DNA sequences.

DNA is often described as the “language of life”, but this analogy can be misleading if interpreted too literally. Natural language and genomic sequences are both discrete symbolic sequences. However, the processes that generate them are fundamentally different. Natural language is produced by humans for communication; it is shaped to transmit information through compact, structured, and semantically dense expressions. Genomic sequence, by contrast, is shaped over evolutionary timescales by mutation, recombination, drift, duplication, transposition, and natural selection [43, 44]. It is not optimized for compact semantic communication. Instead, genomes are highly redundant, only partially constrained, and contain large amounts of background sequence whose functional relevance may be weak, context-dependent, lineage-specific, or currently unknown.

This distinction has direct consequences for pre-training DNA foundation models. In natural language modeling, adding more text usually adds more meaningful linguistic contexts, because most tokens are produced as part of communicative acts. In genomic language modeling, adding more raw sequence can instead introduce large amounts of weakly constrained background, repetitive elements, low-complexity regions, transposon-derived sequence, assembly artifacts, and lineage-specific neutral variation. These regions are not necessarily useless, but they can dominate the training distribution while contributing relatively little to the biological capabilities we want the model to acquire. As a result, pre-training loss alone can be a poor proxy for biological usefulness: a model may obtain lower loss by modeling abundant and locally predictable background patterns, without learning the functional sequence grammar required for sequence recovery, variant interpretation, or regulatory modeling.

### 3.1 A Signal-to-Noise View of Genomic Pre-training

We therefore view genomic pre-training as a signal-to-noise problem, much as modern natural-language data pipelines treat aggressive curation as the key lever for improving training signal density [45]. Let a raw genomic corpus be decomposed into informative regions *S* and background regions *N*, and let *ρ* denote the fraction of informative tokens. Assume that gradients from informative regions have expected alignment *µ* with downstream biological objectives, while gradients from background regions have weaker or less consistent alignment. A simplified effective signal-to-noise ratio for all-sequence training can be written as

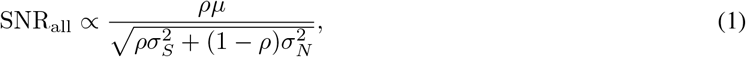

where 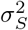 and 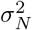 denote the gradient variance contributed by informative and background regions, respectively. When *ρ* is small, even a very large corpus can have a limited effective biological signal, because most optimization steps are dominated by background sequences.

Concretely, we want to selectively retain genomic intervals supported by prior biological evidence, such as genecentric regions and regulatory elements, while dropping or downweighting weakly constrained background. We refer to this as *annotation-aware data curation*. It changes the effective training distribution by enriching regions with a higher prior probability of biological function. Suppose the curation procedure retains a fraction *r* of informative regions and a fraction *ϵ* of background regions. The effective informative fraction after curation becomes

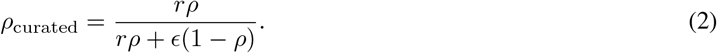

When *ϵ* ≪ *r*, the informative fraction can increase substantially even if some true functional regions are missed. For example, if only 5% of a raw corpus is highly informative, retaining 80% of informative regions while retaining 5% of background regions increases the effective informative fraction from 5% to approximately 46%. This calculation illustrates why a smaller but better curated corpus can be more useful than a much larger corpus dominated by weakly informative sequences.

The same intuition is also consistent with a simple evolutionary search argument. Suppose that a viable biological system requires *K* functional sequence modules, with module lengths *ℓ*_1_, &, *ℓ*_*K*_ and total functional length *ℓ* = ∑_*i*_ *ℓ*_*i*_. Let *p*_*i*_ denote the probability that a random segment of length *ℓ*_*i*_ realizes module *i*, including functionally acceptable variants. If the genome were perfectly compact, with total length *L* = *ℓ*, then all required modules would need to appear in a nearly fixed arrangement, and the probability of such a compact solution scales as

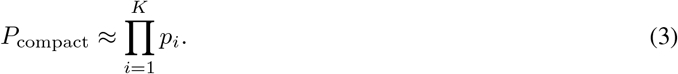

For exact nucleotide targets, this reduces to 4^−*ℓ*^; even for only *ℓ* = 1000 functional bases, the probability is on the order of 10^−602^. This illustrates how restrictive a genome would be if every nucleotide were required to be indispensable.

In contrast, a longer genome with *L* ≫ *ℓ* allows functional modules to be embedded within an arbitrary background sequence. The modules no longer need to form a single compact string; background sequence can appear before, after, or between functional modules. If we keep the order of the *K* modules fixed for simplicity, the number of possible layouts is

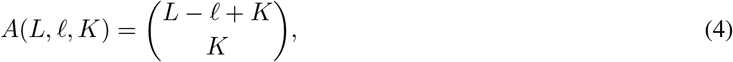

corresponding to the number of ways to distribute the *L* − *ℓ* background bases across the *K* + 1 gaps around the modules. In a rare-event approximation, the probability of obtaining a viable redundant arrangement scales as

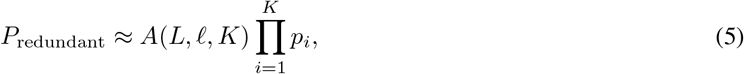

up to probability saturation. Thus, compared with the perfectly compact case, the redundant genome gains a large combinatorial factor *A*(*L, ℓ, K*). For example, with *ℓ* = 1,000, *K* = 100, and *L* = 100,000, this layout factor alone is on the order of 10^240^. Allowing the module order to vary would further multiply the number of valid arrangements by up to *K*!; for *K* = 100, Stirling’s approximation gives 100! ≈ 10^158^, yielding an overall combinatorial factor on the order of 10^398^.

This toy calculation is not intended as a literal model of genome evolution. Real genomes are not random strings, functional elements are degenerate and context-dependent, and evolution proceeds through iterative mutation, selection, duplication, transposition, and drift rather than one-shot sampling. Nevertheless, the argument captures an important asymmetry: a perfectly compact genome in which every base is essential is an extremely restrictive object, whereas sparse functional modules embedded in a longer redundant background occupy a vastly larger space of viable configurations. From this perspective, it is unsurprising that natural genomes are not maximally compact semantic strings, but instead contain functional islands distributed within large amounts of weakly constrained sequence.

Redundancy is also compatible with biological robustness. A genome in which every nucleotide is essential would be extremely fragile: most mutations would have deleterious consequences. By contrast, a genome with distributed functional elements embedded in a larger background can tolerate many neutral or nearly neutral mutations while preserving the modules required for survival and reproduction. This organization provides raw material for evolutionary innovation through duplication, divergence, transposition, and regulatory rewiring. Consistent with this view, eukaryotic genomes contain abundant repetitive elements and transposon-derived sequences, and recent chromosome-engineering experiments in *Arabidopsis thaliana* show that plants can tolerate large-scale karyotype restructuring: CRISPR-Cas-mediated chromosome arm fusions reduced the chromosome number from ten to eight, while the resulting homozygous lines remained fertile and showed no detectable phenotypic or transcriptional differences relative to wild type [46]. Such observations do not imply that background sequences are irrelevant, but they do illustrate that genome organization is often robust, redundant, and only partially constrained at the nucleotide level.

This perspective motivates annotation-aware curation. The goal is not to claim that unannotated or discarded regions are functionless. Rather, under finite compute, pre-training benefits from increasing the density of biological signal in the training distribution. If preserving a small amount of additional useful information requires adding a much larger amount of weakly constrained background, the resulting corpus may become less favorable from a signal-to-noise perspective. Annotation-aware data curation is therefore a practical strategy for improving the effective biological signal seen by the model, while acknowledging that some useful information may still exist outside current annotations.

### 3.2 Carbon Training Corpus

The Carbon training corpus is constructed as a mixture of two annotation-aware data sources (Figure 2). The first source is the GENERator-v2 pre-training data [17]; for brevity, we refer to this corpus as Gener data. It provides the main functional genomic backbone of the Carbon corpus. The second source is OpenGenome2 [9], the pre-training corpus used by Evo2. From OpenGenome2, we use the mRNA-splice, mRNA, and prokaryotic genome components (GTDB and IMG/PR) [47]. We arrive at the final mixture through a series of controlled ablations described below; the resulting composition is summarized at the end of this section in Table 2.

**Figure 1:**
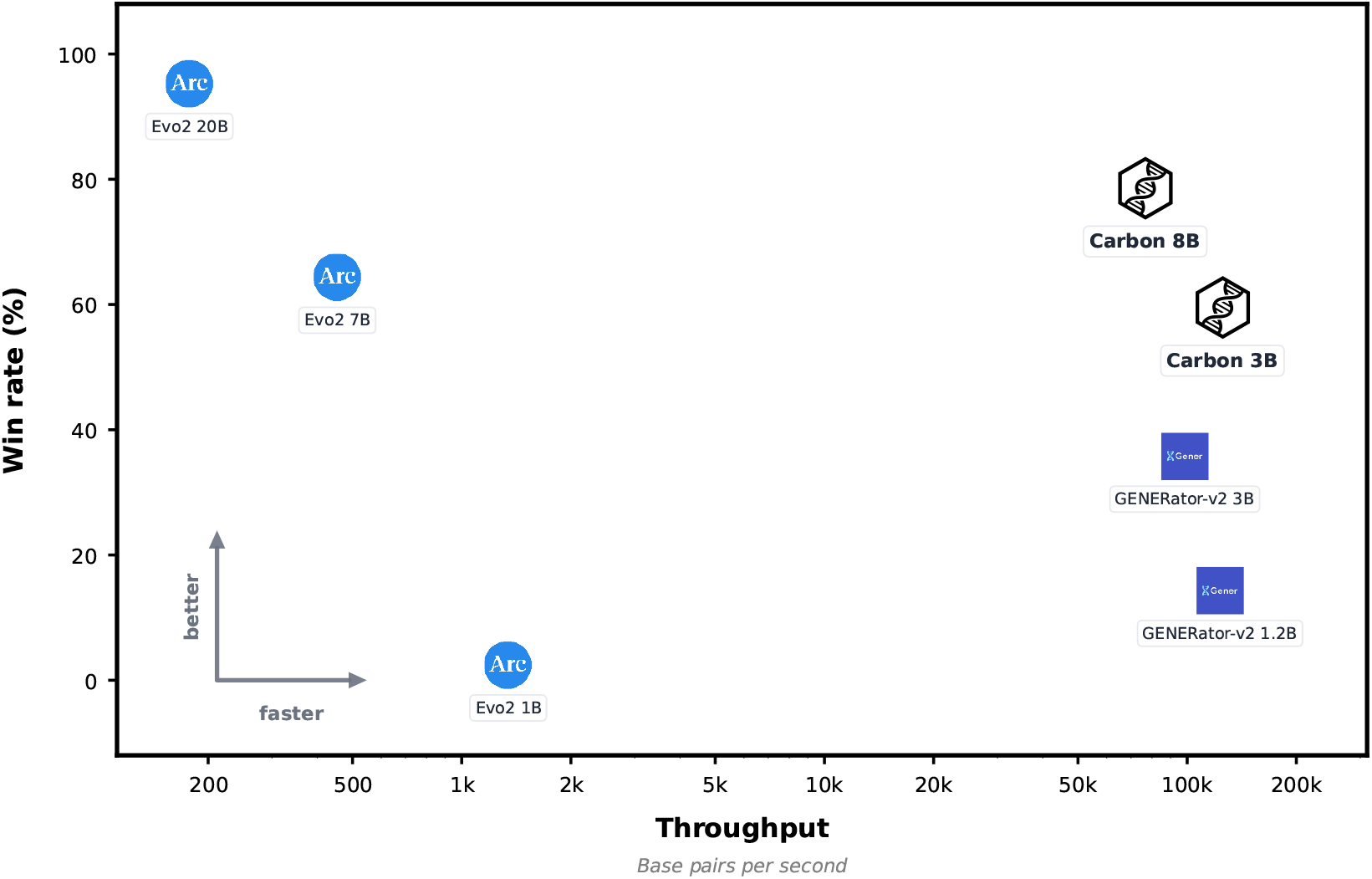
Carbon models obtain comparable or better performance than Evo2 7B, while being over 150 times faster. Win rates computed across seven zero-shot DNA benchmarks. Throughput computed over 256 DNA sequences, with 1080 base pairs for prefill and decode.

**Figure 2:**
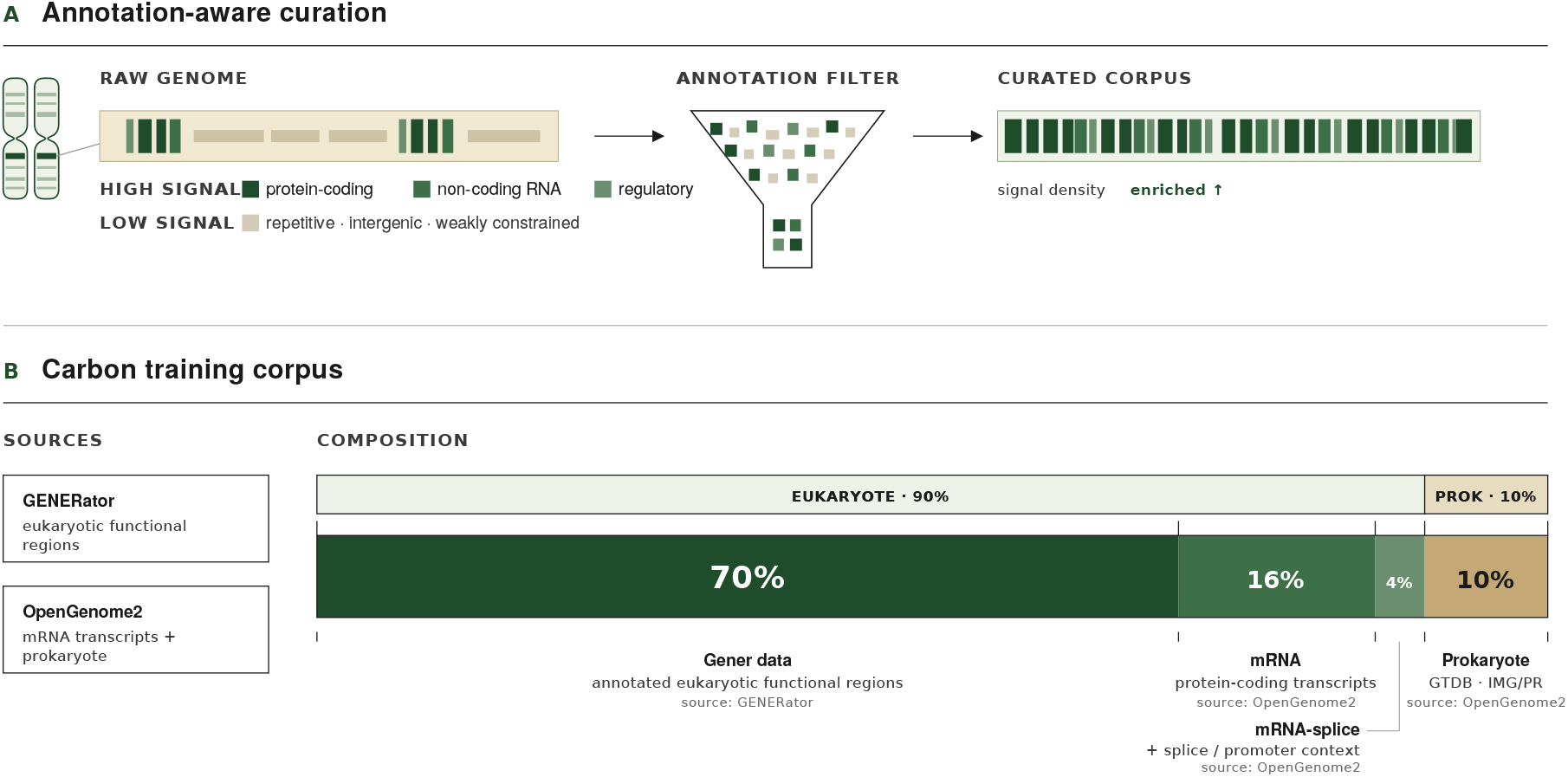
Overview of the Carbon pre-training data: source corpora, annotation-aware curation steps, and final mixture used for Carbon-3B and Carbon-8B.

During base pre-training, Carbon uses a context length of 8,192 tokens. Under 6-mer tokenization, this corresponds to approximately 49k base pairs, which covers many individual functional regions while keeping training efficient enough for 3B/8B-scale models and systematic ablations. The 3B model is later extended to a 32,768-token training context (and a 65,536-token inference context via YaRN [48]) during the long-context phase; details in Section 5.

#### Ablation setup

Prior genomic foundation model work has established that data choices matter: GENERator showed that all-sequence training can achieve lower pre-training loss while producing weaker biological performance, motivating functional sequence training [10]; Evo2 reported that downweighting repetitive regions improves downstream performance [9]. However, many practical data decisions in genomic pre-training are still made without controlled comparisons, making it difficult to tell which choices are actually useful. We therefore test the main choices in our data curation through ablations on a 3B reference model, using the same architecture as the final Carbon-3B (Section 5.1), trained for 50 B tokens at an 8,192-token context length with the 6-mer tokenizer. The ablations cover corpus source, length filtering, sequence construction, and dataset mixing.

All runs are evaluated on four training-free metrics: Sequence Recovery (next-token nucleotide accuracy), BRCA2 (variant-effect prediction AUROC), nucleotide triplet-expansion (perturbation discrimination), and synonymous codon replacement (perturbation discrimination). ClinVar and TraitGym Mendelian are held out from ablation selection and reported only on final models. Full task descriptions are given in Section 6.

#### Choosing the eukaryotic source

We compare two candidate eukaryotic sources by training on each one under matched compute: Gener data, and the OpenGenome2 eukaryote subset. Gener data beats OpenGenome2 on every metric (SR 45.82 vs 43.25, BRCA2 79.22 vs 75.04, with much larger gaps on the perturbation tasks; full curves in Appendix B.1). Gener data also keeps sequences largely as-is, applying only the removal of non-functional regions from individually annotated RefSeq genes, whereas OpenGenome2 layers on additional processing steps. We therefore adopt Gener data as our eukaryotic corpus.

#### Sequence construction

Following the length filter applied in GENERator-v2 training [17], we drop sequences longer than 100 kbp and find that this boosts downstream performance over training on the unfiltered corpus, where the data is dominated by very long, mostly background sequences (Appendix B.3; at 100 kbp the model reaches SR 46.38 / BRCA2 81.16 vs 42.75 / 69.05 with no filter, and triplet-expansion drops from 47.9% to 20.5% without the filter). We also tested a higher 200 kbp threshold, which sits between the two; 100 kbp works best. We also compare concatenated Gener sequences (where consecutive functional regions from the same contig are joined, following GENERator-v2’s Genome Compression Pretraining [17]) against non-concatenated windows at the 8,192-token pretraining context. Non-concatenated windows perform slightly better on every metric (Δ ≈ 0.6–2 pp; Appendix B.2). Unlike GENERator-v2, which used concatenation throughout pretraining (at a 16 k-token context), we therefore adopt non-concatenated windows for base pretraining and defer concatenation to our long-context extension phase (Section 5), where it serves its original purpose of populating long training sequences with biologically informative content.

#### Adding transcript-level data: mRNA vs mRNA-splice

With the eukaryote backbone fixed, we extend coverage beyond DNA-only windows by mixing in transcript-level data from OpenGenome2 [9]. The mRNA subset consists of representative protein-coding transcripts extracted from NCBI GTF annotations and clustered at 90% identity to reduce redundancy; the mRNA-splice subset adds splice and promoter context (Table 2). Both subsets improve every metric over the GENERator-only baseline. 80/20 with mRNA gives the larger gain on SR (+4.5 pp) and triplet-expansion (+12 pp); on BRCA2 and synonymous codon, the two mixes are essentially tied, with differences well under our ~ 2 pp noise floor (Appendix B.4). We therefore allocate the larger share to mRNA (16%) and the smaller to mRNA-splice (4%) in the final mixture.

#### Adding prokaryote coverage

Our primary evaluation focus is on eukaryote tasks, but we want the resulting model to be a good starting point for finetuning on prokaryote applications. We therefore include a small share of prokaryotic genomes (GTDB and IMG/PR, from OpenGenome2) provided this does not hurt eukaryote downstream performance. We find that adding up to 20% prokaryote data leaves eukaryote downstream performance unchanged while improving prokaryote sequence recovery; we adopt 10% to leave more budget for eukaryote and transcript-level data.

#### Metadata conditioning

For a subset of training examples, we prepend metadata tokens describing the species and gene type before the DNA sequence. This provides a weak biological context without changing the autoregressive objective, and gives the model a simple interface for conditional sequence modeling, allowing it to learn how genomic sequence distributions vary across taxonomic and functional categories.

To prevent the model from becoming overly dependent on metadata, we use a mixture of metadata-present and metadata-absent formats: each training sequence is sampled into one of four prompt templates (Table 1). The template mixture serves three purposes: the metadata-free format preserves the standard unconditional DNA language modeling setting and keeps the model usable when no annotation is available; the species-conditioned format encourages the model to represent taxonomic variation and species-specific sequence statistics; and the gene-type-conditioned format encourages it to associate sequence grammar with functional categories such as coding, non-coding, transcript-derived, or splice-related regions. The strategy is analogous in spirit to conditional language modeling, but the metadata is biological rather than linguistic: the goal is not to turn metadata into labels for a supervised classifier, but to expose the model to structured biological context while preserving next-token prediction as the central training objective.

**Table 1:** Metadata template mixture used during Carbon pre-training.

| Sampling fraction | Input template |
| --- | --- |
| 50.0% | <dna>SEQUENCE</dna> |
| 16.7% | <species_type><gene_type><dna>SEQUENCE</dna> |
| 16.7% | <species_type><dna>SEQUENCE</dna> |
| 16.7% | <gene_type><dna>SEQUENCE</dna> |

**Table 2:**
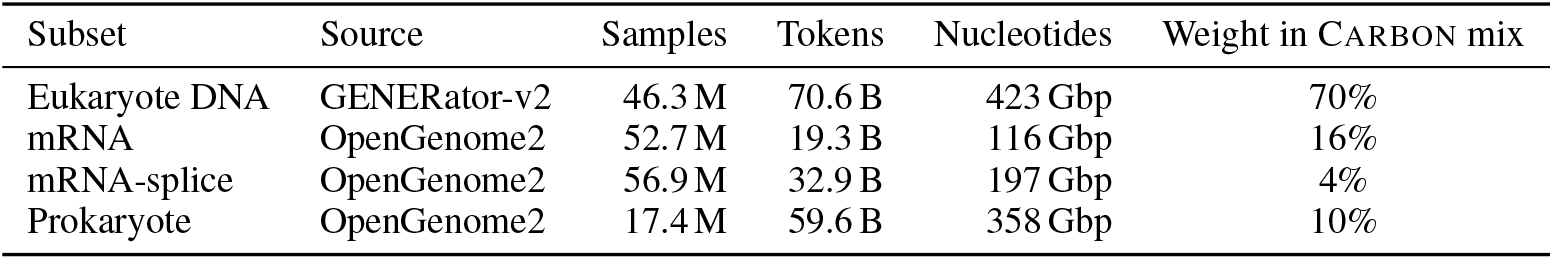
Composition of the Carbon pre-training corpus. Mix fractions correspond to the base pretraining stage (Section 5)

In our ablations, training with tags on 50% of samples leaves overall sequence recovery essentially unchanged. The benefit shows up at inference: supplying a species tag improves sequence recovery on the lowest-resource eukaryote groups, fungi, protozoa, and invertebrates, while leaving high-resource performance unchanged.

#### Data mixing ratios

With each component validated individually, we test several overall mixing ratios across the eukaryote (Gener), transcript-level (mRNA and mRNA-splice from OpenGenome2), and prokaryote (GTDB and IMG/PR from OpenGenome2) components. The combination that performs best on our downstream evaluations is 70% eukaryote, 20% transcript-level (16% mRNA, 4% mRNA-splice), and 10% prokaryote, summarized in Table 2.^1^

This is the base pretraining mixture. We further shift the mixture during the decay phase (upsampling mRNA and prokaryote) and during the long-context extension phase (adding long Gener concatenations and a small share of OpenGenome2 promoters); these stage-specific mixtures are described in Section 5.

### 3.3 Summary

The training data of Carbon is designed around a simple principle: genomic modeling is limited not only by data scale, but by the effective density of biological signal. DNA contains extensive redundancy and background variation, so the raw nucleotide count is an insufficient measure of corpus quality. Annotation-aware curation increases the signal-to-noise ratio of pre-training by enriching functional and transcript-associated regions, while controlled mixture design broadens biological coverage without overwhelming the functional signal.

## 4 Tokenization for Efficient Autoregressive DNA Modeling

After constructing the training corpus, the next key design choice is how to tokenize DNA sequences. Tokenization determines the effective sequence length seen by the model, the computational cost of attention, the granularity of prediction, and the inductive biases imposed on the model. For genomic language models, this choice is especially important because many biologically meaningful tasks require single-nucleotide sensitivity, while many relevant dependencies occur over long genomic spans. A practical tokenizer must therefore balance nucleotide-level fidelity, context coverage, generative stability, and inference efficiency.

In principle, the most direct representation is single-nucleotide (character-level) tokenization, where each base in {*A, C, G, T}* is treated as one token. This is the choice in some recent genomic foundation models such as Evo2 [9]. It preserves maximal input resolution and avoids any ambiguity introduced by grouping nucleotides. However, it is also computationally expensive. For a DNA sequence of length *L*, a single-nucleotide tokenizer produces *L* tokens, whereas a fixed 6-mer tokenizer (which groups every six consecutive nucleotides into a single token drawn from a vocabulary of 4^6^ = 4,096 blocks) produces approximately *L/*6 tokens. For Transformer models, the prefill attention cost scales quadratically with token length, so reducing the token count by a factor of six can reduce the attention cost for the same nucleotide span by up to a factor of 36. Autoregressive generation also benefits directly, since generating the same number of nucleotides requires fewer decoding steps. This efficiency gap is not merely an implementation detail; it determines which models can be evaluated, deployed, and used at scale.

We therefore focus on compressed tokenization. This does not mean that single-nucleotide modeling is unimportant. Models such as Evo2 provide a strong reference point for single-nucleotide genomic modeling. Rather, our goal in Carbon is to investigate whether a compressed tokenizer can retain strong biological capability while substantially improving inference efficiency and effective context coverage. In this setting, the central question becomes: how should multiple nucleotides be grouped into tokens?

We compare two tokenization families: learned byte-pair encoding (BPE) and fixed-length 6-mer tokenization. BPE has been highly successful in natural language processing and forms a core component of many modern LLM tokenizers. Its success relies on a natural assumption: frequently repeated character sequences often correspond to reusable lexical or sublexical units. In natural language, this assumption is well aligned with the existence of words, morphemes, and relatively stable semantic units. In DNA, however, this assumption becomes much less reliable. DNA contains motifs rather than words, and motifs usually do not have sharp boundaries or exact spelling. For example, a canonical motif such as the TATA box represents a family of functionally related sequence patterns rather than a single fixed string. Variants with substitutions, insertions, deletions, length changes, or shifted boundaries may retain similar biological function. Thus, the “vocabulary” of DNA is intrinsically fuzzy: functional units are often degenerate, context-dependent, and only partially specified.

This difference creates a fundamental mismatch between BPE and autoregressive DNA generation. BPE constructs a hierarchical vocabulary in which tokens can overlap semantically and even contain one another as prefixes. For example, if the next nucleotide sequence is `ATCG`, then tokens such as `A, AT, ATC`, and `ATCG` may all be plausible partial continuations under the underlying nucleotide sequence. However, under teacher forcing with a fixed BPE segmentation, only one token is treated as the correct next-token label. The model is therefore penalized for predicting valid prefixes of the target sequence. This ambiguity is minor in masked language modeling, where each masked position is associated with a fixed target token, but it directly interferes with autoregressive next-token prediction. In other words, BPE introduces a learned boundary structure that is useful for natural language but poorly aligned with causal DNA generation.

### Tokenizer ablation

GENERator [10, 17] previously evaluated a broad range of tokenizers for autoregressive DNA modeling: fixed *k*-mer tokenizers for *k* = 1 to 8 and BPE tokenizers with vocabulary sizes from 512 to 8192. Their tokenizer ablation reports average sequence recovery accuracy across six taxonomic groups; across that sweep, neither the highest-resolution single-nucleotide nor the longest-context 8-mer tokenizer was optimal: 6-mer tokenization gave the strongest sequence recovery, while BPE tokenizers underperformed even though they had been successful in masked genomic language models.

Building on this, we run the tokenizer ablations at our reference scale using the same setup as our data ablations (Section 3): a 3B reference model trained for 50 B tokens on the OpenGenome2 eukaryote subset, evaluated on the full training-free suite (Section 6). We compare the fixed 6-mer tokenizer against two BPE tokenizers trained on the same eukaryote data at LLM-scale vocabulary sizes of 32 k and 50 k tokens. As predicted by the next-token mismatch argument above, 6-mer roughly doubles SR over BPE-32k and BPE-50k (43.25 vs 21.68 / 23.48) and improves BRCA2 AUROC by ~ 10 pp (75.04 vs 63.40 / 66.51) at matched compute (Figure 3). The two BPE variants score higher on nucleotide triplet-expansion, but this is a tokenization artefact: BPE merges the (CAG)_10_ patch into two long tokens that look anomalous out of context, while 6-mer’s frame-aligned grid splits the patch into ordinary tokens that hide inside the lattice. Sequence Recovery, the metric most directly tied to next-token DNA modelling, clearly favours 6-mer, as does the variant effect prediction task BRCA2.

**Figure 3:**
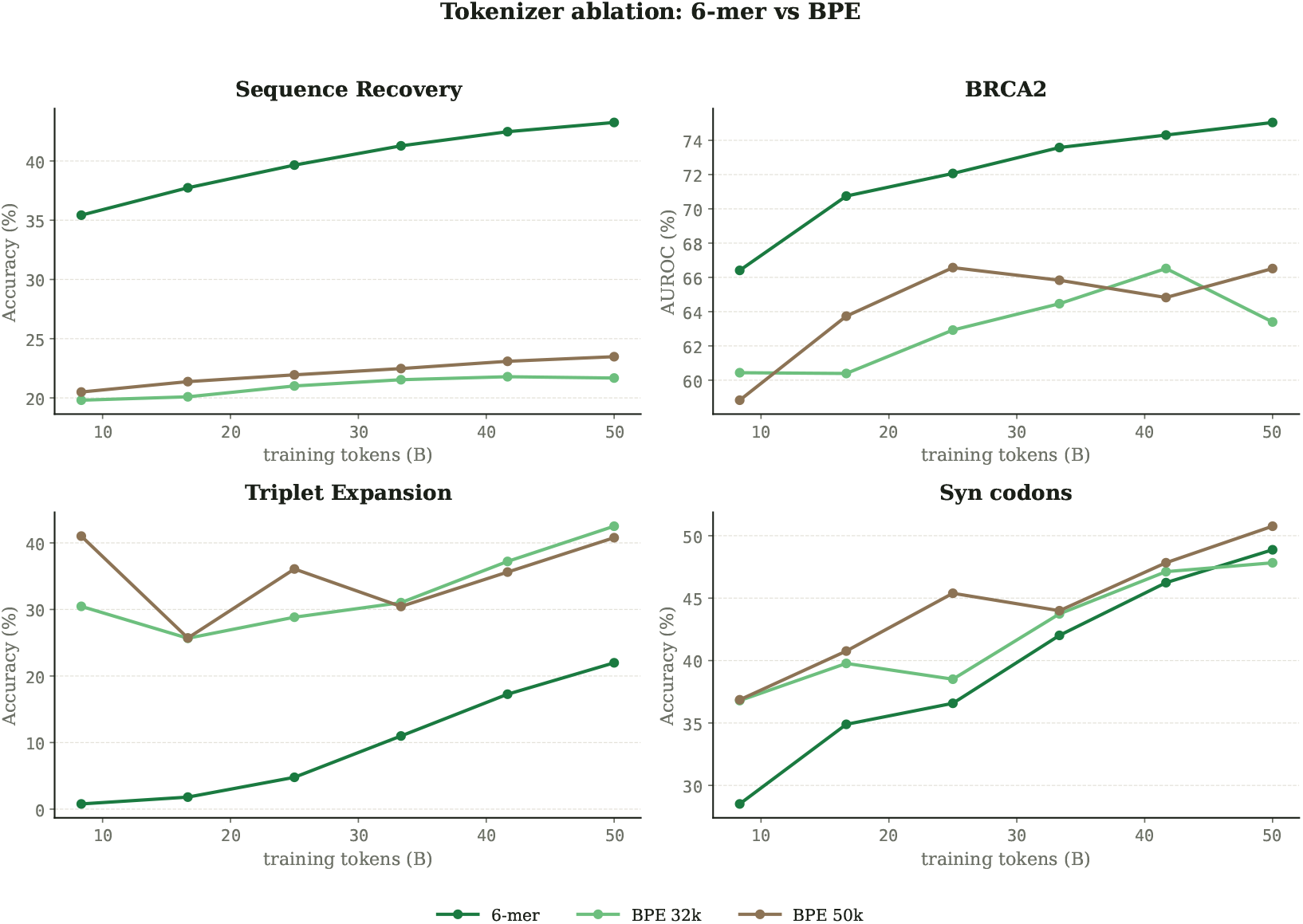
Tokenizer ablation at matched compute (3B, 50 B tokens, OpenGenome2 eukaryote windows). The fixed 6-mer tokenizer roughly doubles sequence recovery over BPE at 32 k and 50 k vocabulary sizes and improves BRCA2 AUROC by ~10 pp.

We choose 6-mer tokenization for Carbon not because DNA is biologically organized in units of six nucleotides. Rather, 6-mer is a simple, deterministic, and largely neutral compression scheme. It does not attempt to discover word-like boundaries in DNA, nor does it impose a learned vocabulary of variable-length motifs. Instead, it maps every length-six nucleotide block into one of 4^6^ = 4,096 possible tokens:

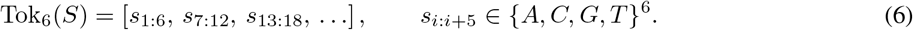

Each 6-mer token is then embedded into a high-dimensional continuous space by the model. This gives the model enough representational capacity to learn local sequence patterns while reducing the sequence length by a factor of six. Conceptually, this can be viewed as processing and predicting DNA in fixed-size blocks: compared with a singlenucleotide model, the model reads six nucleotides at a time and predicts six nucleotides at a time, while representing each block through a learned embedding.

This blockwise view also clarifies why 6-mer tokenization is compatible with nucleotide-level reasoning. Although the model operates over a 4,096-way token vocabulary, each token has an explicit internal structure composed of six nucleotides. Later in training, this structure can be exposed through nucleotide-aware supervision, such as Factorized Nucleotide Supervision (FNS), which marginalizes token-level probabilities into nucleotide-level probabilities. Thus, 6-mer tokenization provides computational compression without making nucleotide-level inference impossible. It trades input granularity for efficiency, but the biological resolution can be partially recovered through the training objective and inference procedure.

The choice of *k* = 6 reflects an empirical balance. Smaller *k* values preserve finer token-level resolution but provide less context coverage under a fixed token budget. Larger *k* values further improve compression but make each token increasingly coarse and enlarge the vocabulary as 4^*k*^, making the prediction problem sparser. A 6-mer vocabulary of 4,096 tokens is large enough to encode rich local sequence patterns, yet small enough to remain stable for autoregressive training. Under an 8k-token context window, this tokenizer allows Carbon to process up to approximately 8,192 × 6 = 49,152 base pairs, which provides substantial genomic context while preserving practical inference cost.

Importantly, we do not claim that 6-mer is the final or universally optimal tokenizer for DNA. It is best understood as a conservative and effective choice: it introduces minimal biological assumptions, avoids the hierarchical ambiguity of BPE, and provides a large efficiency gain over single-nucleotide tokenization. More sophisticated tokenizers may eventually learn better representations of motifs, regulatory grammar, or evolutionary conservation. However, such tokenizers must be evaluated under the objective for which they are intended. A tokenizer that works well for masked language modeling or natural language processing does not necessarily work well for autoregressive DNA generation. For Carbon, fixed 6-mer tokenization provides the right combination of simplicity, efficiency, stability, and empirical performance.

### Hybrid tokenization for joint DNA and natural-language modeling

6-mer tokenization is well-matched to a pure-DNA generative model, but it would impose constraints in settings where the model must also process natural language: e.g. a single model trained jointly on DNA sequences and biomedical text, or DNA segments interleaved with natural-language descriptions of experiments and annotations. To leave that door open, Carbon’s tokenizer is hybrid by construction: we augment the BPE vocabulary of the Qwen3 tokenizer [49] with 4,096 fixed 6-mer DNA tokens and the structural `<dna> / </dna>` delimiters (Section 5.1). On pure DNA inputs, the regime trained and evaluated in this report, the tokenizer behaves exactly like a fixed 6-mer tokenizer, because only the 6-mer part of the vocabulary is activated inside the `<dna> & </dna>` block. Actually exercising the BPE part of the vocabulary for joint DNA + English training is left for future work.

In summary, the tokenizer design of Carbon follows the same principle as its data construction: avoid importing assumptions from natural language modeling without biological and empirical validation. BPE is powerful for human language because human language has relatively stable lexical units. DNA sequence does not provide the same kind of word boundary. By using fixed 6-mer tokenization, Carbon adopts a neutral compression strategy that reduces token length by six-fold, improves inference efficiency and context coverage, and preserves a well-defined autoregressive objective.

## 5 Training

This section describes how Carbon is built and trained. We first specify the architecture and then walk through the training recipe: the choice of training objective for 6-mer DNA modeling, the pretraining stages, and the long-context extension phase used to reach the final training context length of both Carbon-3B and Carbon-8B.

### 5.1 Model Architecture

Carbon uses a standard decoder-only Transformer [2] architecture based on Llama 3 [50]. We release two model sizes: Carbon-3B and Carbon-8B, sharing the same backbone family and differing only in width, depth, and number of attention groups.

Each model is a pre-norm causal Transformer with RMSNorm [51], SwiGLU [52] feed-forward blocks, rotary position embeddings (RoPE; Su et al. 53), and grouped-query attention (GQA; Ainslie et al. 54). The base frequency *θ* = 500,000 (from Llama 3 [50]) is used during pretraining and rescaled to *θ* = 5,000,000 in the long-context extension phase. The vocabulary extends the Qwen3 BPE tokenizer [49] with 4,096 DNA 6-mer tokens, three DNA structural tokens (`<dna>, </dna>, <oov>`), and padding tokens for hardware alignment, for a total vocabulary size of 155,776. Input and output token embeddings are tied; this regularises the large DNA sub-vocabulary whose tokens are seen infrequently relative to BPE text tokens. Notably, Carbon-8B shares its layer count, hidden size, feed-forward dimension, and attention configuration exactly with Llama 3-8B, making it a useful reference point for isolating the contribution of data and objective design from architectural differences. The hyperparameters are summarized in Table 3.

**Table 3:**
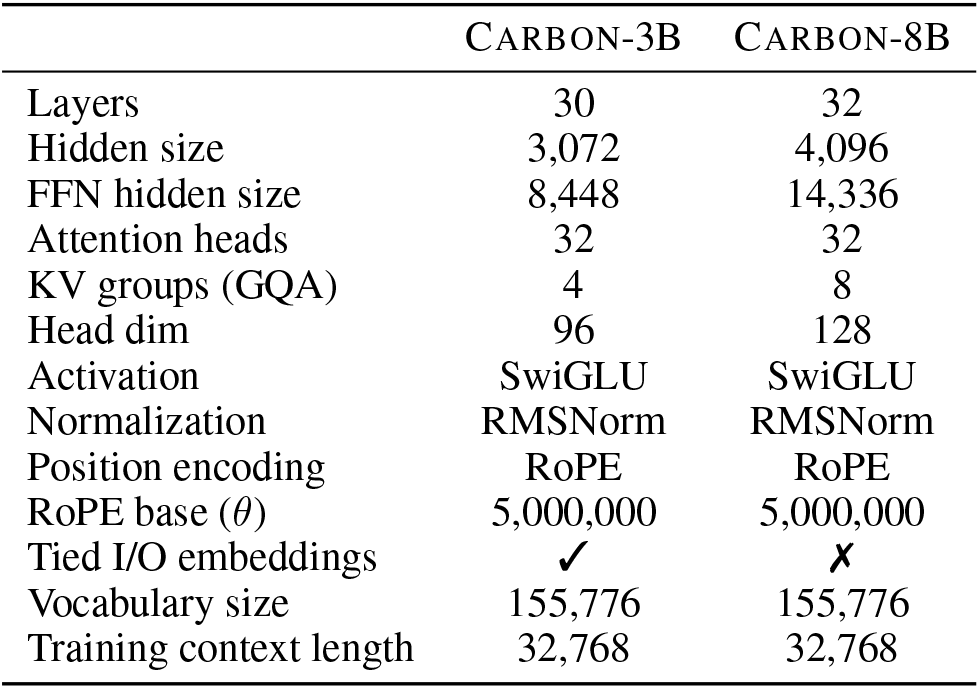
Architecture hyperparameters of Carbon-3B and Carbon-8B.

Both models pretrain at an 8,192-token context length (roughly 49 kbp under 6-mer tokenization) and are then extended to 32,768 tokens (~ 197 kbp) in a training-time long-context phase. At inference, YaRN [48] further extends Carbon-3B to 65,536 tokens (~ 393 kbp, 2*×*) and Carbon-8B to 131,072 tokens (~ 786 kbp, 4*×*); the larger YaRN factor for Carbon-8B reflects the greater extrapolation headroom afforded by its larger capacity (details below).

This configuration is intentionally close to widely used open-weight language model families. The aim is to keep the architectural surface neutral, so that any performance characteristics observed on genomic tasks reflect the modeling choices specific to Carbon: the data composition described in Section 3, the 6-mer tokenization described in Section 4, and the training recipe described below. Architectural improvements such as attention variants and hybrid state-space layers are orthogonal to the questions investigated in this work and are left for future study.

### 5.2 Training Objective

Beyond the architecture, the next key design choice is the training objective. For autoregressive language models, the standard objective is next-token cross-entropy (CE). CE is simple, effective, and remains the natural starting point for generative DNA modeling. However, when combined with non-overlapping 6-mer tokenization, CE treats each 6-mer as an atomic class among 4^6^ = 4,096 possibilities. As a result, supervision is applied only at the full-token level and does not reflect partial nucleotide agreement between nearby sequence predictions. Such exact-token supervision is useful for learning joint nucleotide dependencies within a 6-mer, but it can become too sharp for DNA, where many functional elements are degenerate, context-dependent, and naturally variable at the nucleotide level.

In preliminary CE-only training runs, we observed a recurring instability that we refer to as the *loss staircase* (Figure S1). Rather than showing only transient spikes, the smoothed training loss could jump upward and remain elevated for an extended period. This instability was accompanied by a widening gap between BF16 [55] and FP32 sequence-recovery accuracy: after the staircase, FP32 generation could remain stable or continue improving, while BF16 generation degraded substantially. This suggests that late-stage CE training can push the model into a numerically fragile regime, where correct generation depends on fine probability distinctions among many plausible 6-mer candidates.

To address this issue, Carbon uses Factorized Nucleotide Supervision (FNS) [17]. FNS preserves the 6-mer output space, but computes the loss through nucleotide-level marginals induced by the 6-mer distribution. For each position *j* ∈ {1, …, 6} and nucleotide *b* ∈ {*A, C, G, T*}, the marginal probability is

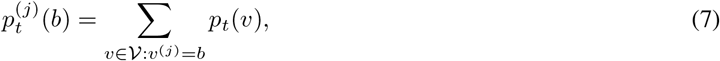

and the FNS loss for target 6-mer *y*_*t*_ is

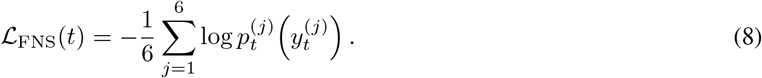

This objective gives structured partial credit to near-miss tokens: a 6-mer that matches five positions contributes probability mass to five correct nucleotide marginals, rather than being treated identically to a completely mismatched token. This is important because DNA sequence is not always an all-or-nothing object. Many regulatory motifs are degenerate and context-dependent; for example, a TATA-like element may remain functional even when one nucleotide differs from a canonical pattern. Exact 6-mer CE cannot express this graded similarity: it treats every non-target 6-mer as equally wrong, which can make late-stage optimization overly sharp and numerically fragile. FNS addresses this problem by replacing binary exact-token supervision with a smoother nucleotide-aware signal, while still preserving the same ideal optimum: the exact target 6-mer remains the only token that satisfies all six nucleotide constraints simultaneously.

We do not, however, train with FNS from scratch. In our experiments, FNS-from-scratch removes the loss staircase but leads to weaker performance under the same training budget. The reason is that FNS relies on a conditional-independence approximation among the six nucleotide positions within a 6-mer, and this approximation is not always appropriate. In coding regions, splice sites, and other sharply constrained contexts, a single-nucleotide change can alter an amino acid, introduce a premature stop codon, or disrupt a required motif. Early CE training is therefore valuable because its exact-token objective efficiently teaches the joint distribution of nucleotides within each 6-mer. The problem is that, after much of this joint structure has been learned, continued CE optimization can become too rigid and enter the loss-staircase regime. The staged CE-to-FNS schedule follows this division of labor: CE first learns the joint 6-mer structure, and FNS then provides a smoother nucleotide-level refinement objective that stabilizes late-stage training, improves BF16 robustness, and allows the model to continue improving. Based on this observation, Carbon adopts a staged CE-to-FNS objective schedule:

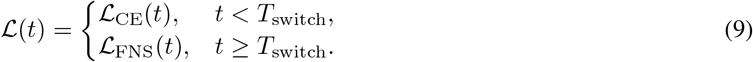

At the switch point, we also reduce the learning rate. The goal is to first let CE teach the model a strong joint-token prior, and then use FNS to smooth late-stage optimization, recover nucleotide-level supervision, and improve robustness under BF16 inference. Training-free sequence recovery is used throughout pre-training as the primary monitoring signal for this transition, because it is tokenizer-agnostic, genuinely generative, non-saturating, and sensitive to both capability growth and numerical instability.

#### FNS for base-pair-level inference

The same nucleotide marginalization used by FNS also provides a practical inference-time mechanism for base-pair-level generation and scoring. For generations, instead of selecting a 6-mer token directly from the full 4,096-way distribution, we marginalize the token distribution into six 4-way nucleotide distributions, choose or sample one base at each position, and then recombine the six bases into the corresponding 6-mer token. This allows generation to operate at nucleotide resolution rather than being determined by the single most likely coarse token.

The same idea also applies to sequence scoring. Given a DNA sequence, the model can assign a probability to each base by marginalizing over all 6-mer tokens that contain that base at the corresponding within-token position. This produces nucleotide-level likelihoods from a coarse-token model and is the key mechanism by which FNS reconciles 6-mer tokenization with single-nucleotide resolution. In our evaluations, FNS-based inference improves sequence recovery, variant effect prediction, and perturbation scoring compared with direct token-level generation or token-level likelihood scoring, at the cost of roughly 35% additional inference time. Detailed derivations, gradient analysis, switch-timing discussion, base-pair-level decoding and scoring procedures, and comparisons to label smoothing, multi-token prediction, and blockwise attention are provided in Appendix A.

### 5.3 Pretraining

Carbon-3B and Carbon-8B share the same training recipe in broad terms: CE pretraining, an FNS phase, and a long-context extension phase. The difference is that Carbon-3B includes a dedicated decay phase with an updated data mixture, whereas Carbon-8B folds the cosine decay into a single long FNS phase on the base mixture due to logistic constraints. Both models then undergo the same long-context extension. The sections below describe each training stage.

Both Carbon-3B and Carbon-8B pretrain at an 8,192-token context length (49 kbp) on the annotation-aware mixture introduced in Section 3 (70% Gener data with metadata, 16% mRNA, 4% mRNA-splice, 10% prokaryote) and target roughly 1 T tokens of pretraining. We ablated the pretraining context length and found that 4 k, 8 k, and 16 k tokens give similar downstream performance, so we adopt 8 k as a balance between efficient training (cheaper than 16 k) and tractable long-context extension later (a shorter jump than starting from 4 k).

Pretraining proceeds in two phases. In the first 100 B tokens, the loss is plain cross-entropy on the 6-mer vocabulary, with a Warmup-Stable-Decay (WSD) learning-rate schedule warmed up over 2,000 iterations to a peak of 3 × 10^−4^. After this CE phase, we switch to the FNS objective at a lower peak learning rate of 5 × 10^−5^, following the dynamics described above for the CE-to-FNS switch.

The FNS stable phase differs across the two models:

- Carbon-3B continues on the base mixture under FNS until the end of the stable phase at 800 B tokens; the mixture is then shifted in the decay phase (below).
- Carbon-8B continues on the base mixture under FNS for the entire remaining 900 B tokens, with a single cosine decay from 5 × 10^−5^ to 5 × 10^−6^ covering the whole span. The model then undergoes the same long-context extension as Carbon-3B (below).

The switch to FNS at 100 B tokens is motivated by the loss staircase observed when CE training is pushed further (Figure S1). After the switch, training remains stable through the rest of pretraining for both models, with no recurrence of the staircase (Appendix C).

#### Decay

During Carbon-3B’s decay phase, we update the training mixture: 50% Gener data, 25% mRNA, 10% mRNA-splice, 15% prokaryote, upsampling mRNA and prokaryote relative to the base 70/16/4/10 mixture. We find that this new mixture slightly improves downstream performance on our evaluation suite.

#### Long-Context Extension

Both Carbon-3B and Carbon-8B undergo a training-time long-context extension phase that lifts the training context length from 8,192 to 32,768 tokens. The RoPE base frequency *θ* is rescaled from 5 × 10^5^ to 5 × 10^6^. We train for 50 B tokens under FNS; the learning rate is warmed up over 2,000 iterations to a peak of 3 × 10^−5^ and cosine-decayed to a floor of 3 × 10^−6^. Throughout training, the loss is computed only on 6-mer DNA token positions; the structural tags (`<dna>, </dna>`) are masked and contribute to neither the CE nor the FNS objective.

The data mixture in this phase builds on the decay-phase mixture with two additions designed to expose long-range structure and improve downstream performance: 13.8% long-context Gener concatenations, formed by joining consecutive functional regions from the same contig in the Gener data, and 1.2% promoter sequences from the OpenGenome2 EPDnew subset [9]. The base Gener share is reduced to 35% to make room for these new components, while mRNA (25%), mRNA-splice (10%), and prokaryote (15%) shares match the decay-phase composition.

At inference, both models are further extended with YaRN [48] from the 32,768-token training context: Carbon-3B to 65,536 tokens (393 kbp, 2*×*), and Carbon-8B to 131,072 tokens (786 kbp, 4*×*). In our long-context retrieval evaluation (Section 6), Carbon-3B holds up cleanly at 2× but loses accuracy when pushed to 4*×*, while Carbon-8B extrapolates to 4× without collapse.

### 5.4 Summary

The Carbon training recipe combines multi-stage pretraining on the annotation-aware mixture, an objective switch that stabilizes late-stage 6-mer learning, and a long-context extension phase.

As for the training objective, it is designed to balance three requirements: learning joint 6-mer dependencies, preserving nucleotide-level biological structure, and maintaining stable, efficient inference. Standard CE is effective in early training because it teaches the joint distribution of 6-mer tokens, but it can become too sharp in late training, producing loss staircases and increased BF16 sensitivity. FNS addresses this issue by replacing exact-token supervision with nucleotide-level marginal supervision, providing structured partial credit and smoother gradients. However, FNS relies on a conditional-independence approximation and operates on block-level hidden states, so it is suboptimal when used from scratch.

The resulting strategy is CE-to-FNS switching. Carbon first learns with CE, then switches to FNS with a reduced learning rate when CE dynamics approach instability. Sequence recovery is used as the primary monitoring signal because it is tokenizer-agnostic, generative, non-saturating, and sensitive to both capability growth and numerical robustness. This objective schedule removes the loss staircase, reduces the BF16/FP32 performance gap, and allows the model to continue improving after pure CE training becomes unstable. The final training recipe, therefore, combines annotation-aware data curation, fixed 6-mer tokenization, and staged CE-to-FNS optimization.

## 6 Training-Free Evaluation Framework

As genomic foundation models continue to scale, evaluation itself becomes a practical bottleneck. Full fine-tuning is expensive, implementation-dependent, and difficult to apply uniformly to large models with different architectures, tokenizers, context lengths, and inference costs. For this reason, we focus primarily on training-free evaluation in this report. By training-free, we refer to protocols that evaluate frozen pre-trained models without task-specific parameter updates. This makes the evaluation suite suitable both for comparing large model families under controlled settings and for tracking model capability during pre-training.

The suite includes biological benchmarks from the literature: ClinVar coding and non-coding variant-effect prediction [19, 20], BRCA2 saturation mutagenesis [21], and TraitGym Mendelian non-coding variants [22]. We complement these with controlled diagnostic probes for autoregressive sequence generation, sequence-level perturbation sensitivity, and long-context retrieval. Together, these tasks support model development by tracking pre-training progress, comparing tokenizers and model scales. These evaluations complement, but do not replace, more biologically specific benchmarks and application-oriented studies for assessing practical biological utility.

We evaluate Carbon along eight training-free tasks across four capability axes: generation (Sequence Recovery), variant effect prediction (ClinVar coding and non-coding, BRCA2, TraitGym Mendelian), sequence-level perturbation (nucleotide triplet-expansion and synonymous codon replacement), and long-context retrieval (Genomic-NIAH). The first two axes probe nucleotide-level generation and likelihood calibration. The third axis covers controlled sequence-level perturbation tasks that test whether the model assigns higher compatibility to natural coding sequences than to synthetic alternatives; the fourth tests long-range retrieval through Genomic-NIAH. Together, these tasks provide a lightweight but informative framework for evaluating large generative DNA models under reproducible inference settings. Table 4 summarizes the suite; the evaluation code is released alongside the models.^2^

**Table 4:** Training-free evaluation suite. The 8 selected tasks span four capability axes.

| Task | Metric | Organisms | Sequence type | # samples | Source |
| --- | --- | --- | --- | --- | --- |
| *Generation* |  |  |  |  |  |
| Sequence Recovery | next-token acc. (%) | 6 groups<sup>a</sup> | mixed | 30,000 | GenerTeam [10] |
| *VEP* |  |  |  |  |  |
| ClinVar coding | AUROC | human | coding | 39,473 | GPN-MSA [20] |
| ClinVar non-coding | AUROC | human | non-coding | 15,258 | Ours |
| BRCA2 | AUROC | human | coding | 6,840 | Huang et al. [21] |
| TraitGym Mendelian | AUPRC (by-chrom) | human | non-coding | 3,380 | TraitGym [22] |
| *Perturbation* |  |  |  |  |  |
| Triplet-expansion | discrimination acc. (%) | human | coding | 20,000 | Ours |
| Codon replacement | discrimination acc. (%) | human + mouse | coding | 20,000 | Ours |
| *Long-context retrieval* |  |  |  |  |  |
| Genome-NIAH | exact-match acc. (%) | 4 kingdoms<sup>b</sup> | mixed | 12,000 | Ours |

<sup>a</sup> Fungi, plants, protozoa, invertebrates, mammals, other vertebrates. <sup>b</sup> Animals, fungi, plants, protists.

### 6.1 Evaluation Protocol and Design Principles

Training-free evaluation serves two roles in this report. First, it enables scalable comparison across large frozen models. Models such as Carbon-3B, Carbon-8B and Evo2-7B differ substantially in parameter count, architecture, tokenizer, maximum supported context length, memory footprint, and generation speed. Fine-tuning all models under identical settings would be computationally expensive and difficult to reproduce. Training-free tasks instead evaluate the pre-trained sequence prior directly, reducing confounding factors introduced by task-specific optimization.

Second, training-free evaluation provides useful signals during model development. Raw training loss is not always comparable across tokenizers: a single-nucleotide model predicts one of four bases at each step, a 6-mer model predicts one of 4^6^ = 4,096 tokens, and a BPE model uses a learned variable-length vocabulary. Moreover, genomic pre-training loss can be affected by repetitive, low-complexity, or weakly constrained sequences. A model can reduce loss by modeling abundant background patterns without necessarily improving the biological capabilities of interest. A useful training-free probe should therefore be tokenizer-agnostic, inexpensive enough to run repeatedly, sensitive to model improvement, difficult enough to avoid early saturation, and interpretable across model families.

Sequence recovery satisfies these requirements particularly well and is used as our primary training-time capability monitor. Variant effect prediction complements it by probing nucleotide-level likelihoods, the perturbation tasks evaluate sequence-level discrimination under controlled modifications, and Genomic-NIAH measures long-range retrieval. Together, these tasks provide a practical evaluation layer before moving to more expensive downstream fine-tuning, application-specific benchmarking, or experimental validation.

### 6.2 Selecting Training-Free Benchmarks

Beyond the general requirements above, we additionally require each candidate task to provide a stable and reliable signal during pre-training. We adopt the four “early-signal” criteria of Penedo et al. [18]:

- **Monotonicity**: performance should improve as training progresses.
- **Low noise**: relative performance differences between models should reflect inherently better training, not evaluation noise.
- **Non-random performance early in training**: tasks reflecting capabilities acquired only late in training cannot meaningfully differentiate experiments at a small scale.
- **Ordering consistency**: model rankings remain stable throughout training.

We test these criteria with intermediate-checkpoint curves for two 3B 6-mer runs trained on Gener data (under identical setups with different seeds) and a 3B 6-mer run trained on the OpenGenome2 eukaryote subset (Figure 4). The selected suite shows smooth monotone improvement, above-random scores from early in training, and consistent ranking between the two data sources throughout training. The two-seed comparison gives a practical reference for how small a metric difference still represents a real signal: end-of-training metrics agree to within ~0.2 pp on SR and synonymous codon and within ~1.5 pp on BRCA2 and nucleotide triplet-expansion.

**Figure 4:**
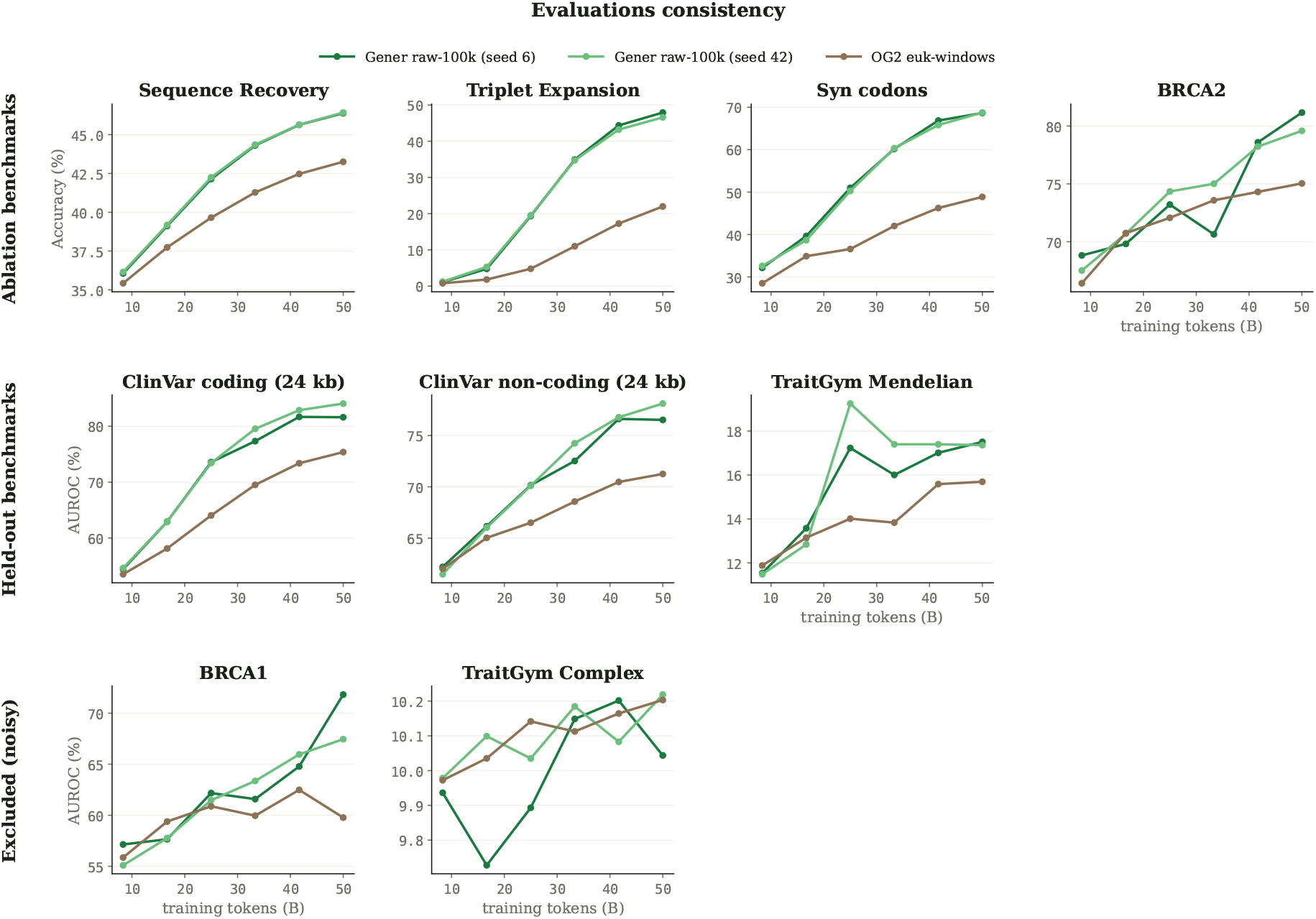
Validating the training-free evaluation suite. Two 3B 6-mer runs trained on Gener data (with different seeds) and one 3B 6-mer run trained on the OpenGenome2 eukaryote subset. The tasks we keep show two-seed runs tracking closely, monotonic improvement across checkpoints, and consistent Gener-over-OpenGenome2 data ranking throughout training.

ClinVar and BRCA2 show a slightly noisier behaviour, with some small gaps between the two 6-mer seeds. We nonetheless retain them in the evaluation suite because they are widely used for variant effect prediction in the genomic foundation-model literature. We only report ClinVar performance on the final models alongside TraitGym Mendelian, rather than using it to choose between ablation variants. We also tested BRCA1 and TraitGym Complex as candidate tasks. BRCA1 was noisier than BRCA2, likely in part because it contains fewer variants (~ 3.9k versus ~6.8k), while TraitGym Complex remained near random for all evaluated models, including Evo2-7B; we therefore drop both from the main suite. The four ablation metrics we use for model selection during development are Sequence Recovery, BRCA2, nucleotide triplet-expansion, and synonymous codon replacement; ClinVar and TraitGym Mendelian are held out.

### 6.3 Nucleotide-Level Prediction Tasks

We first evaluate models on nucleotide-level prediction tasks. These tasks ask whether a frozen pre-trained model can assign meaningful probabilities to individual bases, either through autoregressive generation or by comparing the likelihoods of nucleotide alternatives. They are especially important for genomic foundation models because many biological questions, including variant interpretation, splice-site analysis, codon-level reasoning, and regulatory mutation scoring, depend on single-nucleotide resolution.

#### 6.3.1 Sequence Recovery

Sequence recovery [1, 10] is our primary intrinsic generative evaluation. Given a DNA prompt of fixed length *L*, the model is asked to generate the next *M* nucleotides. The generated continuation is compared with the ground-truth continuation at nucleotide resolution:

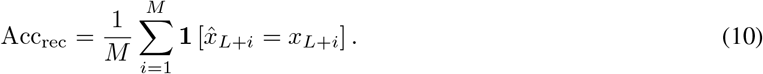

This task plays a role analogous to perplexity tracking or pre-training loss comparison in natural language modeling, but is better suited to genomic models with different tokenization schemes. Because both the input and output lengths are specified in base pairs rather than tokens, sequence recovery can compare single-nucleotide, BPE, and 6-mer models under the same sequence-level setting. This property makes it especially useful for tokenizer ablations; our comparison between BPE and 6-mer tokenization is therefore based primarily on sequence recovery rather than raw training loss.

Sequence recovery is also a genuinely generative task. It evaluates whether a model can produce accurate DNA continuations under autoregressive decoding, rather than only whether it can provide useful representations for a downstream classifier. In our experiments, sequence recovery improves steadily during training, remains non-saturating over a broad range of model scales, and reflects expected trends with model size and training progress. For these reasons, it is our first reference metric for monitoring whether pre-training is proceeding successfully.

#### 6.3.2 Variant Effect Prediction

Variant effect prediction [19, 20] evaluates whether a model assigns biologically meaningful probabilities to singlenucleotide variants. Given a reference sequence *x* and a single-nucleotide variant at position *i*, with reference allele *r* and alternative allele *a*, we compute a log-likelihood ratio:

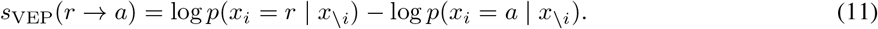

A larger score indicates a stronger model preference for the reference allele over the alternative allele.

For 6-mer models, nucleotide probabilities are obtained by marginalizing over the 6-mer token distribution. If position *i* corresponds to the *j*-th nucleotide inside a 6-mer token, then

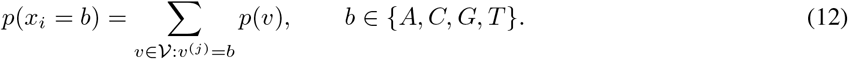

This converts the 4,096-way token distribution into a 4-way nucleotide distribution at the variant position, allowing 6-mer models to perform single-nucleotide scoring without changing the architecture or tokenizer.

Variant effect prediction complements sequence recovery. Sequence recovery measures whether the model can generate the correct continuation, whereas VEP measures whether the model’s probability distribution is calibrated enough to distinguish reference and alternative alleles. Together, these two tasks probe both nucleotide-level generative accuracy and nucleotide-level probabilistic reasoning. Because neither task requires supervised fine-tuning, they provide direct evidence of whether pre-training alone has induced a useful sequence prior at base-level resolution.

##### ClinVar (coding subset)

The coding subset comprises 39,473 exonic protein-coding variants from the GPN-MSA evaluation [20, 19], labelled as pathogenic or benign. We follow the right-end next-token scoring protocol of GEN-ERator [10]: each variant is placed at the right edge of a 24 kb upstream context, and the variant score is the log-ratio of the reference and alternate next-token probabilities. We report AUROC and AUPRC.

##### ClinVar (non-coding subset)

Existing ClinVar dataset from GPN-MSA is heavily skewed toward coding variants: the split contains only 10 pathogenic non-coding variants. We construct a separate balanced non-coding subset directly from the raw ClinVar VCF, retaining single-nucleotide variants on chromosomes 1–22, X, and Y, restricted to reviewed entries with a clean non-coding consequence annotation. The resulting benchmark contains 15,258 variants (7,629 benign / 7,629 pathogenic) spanning intronic and 5^*′*^/3^*′*^ UTR regions. We use the same right-end next-token protocol as the coding subset at 24 kb context and report AUROC and AUPRC.

##### BRCA2 saturation mutagenesis

We use the BRCA2 saturation mutagenesis assay of Huang et al. [21], which provides functional measurements for 6,840 single-nucleotide variants. We treat the task as binary classification (functional vs loss-of-function) and adopt the centered 8 kb-window protocol of Evo2 [9]: each variant is placed at the center of an 8 kb context, and the score is the full-sequence log-likelihood difference between reference and alternate alleles. We report AUROC.

##### TraitGym Mendelian (non-coding)

TraitGym [22] is a recent benchmark suite for variant effect prediction on disease-associated non-coding variants. We use the Mendelian split, which focuses on rare large-effect variants annotated to known Mendelian disease loci, and adopt the centered 8 kb-window scoring used by both the TraitGym authors [22] and Evo2 [9], with reverse-complement averaging of the log-likelihood delta. Following the TraitGym protocol, we report AUPRC under chromosome-stratified evaluation (“by-chrom”).

### 6.4 Sequence-Level Perturbation Tasks

Sequence recovery and VEP focus on nucleotide-level prediction. We additionally use controlled sequence-level perturbation tasks to compare model scores of natural sequences and synthetic alternatives. These tasks are intentionally label-free and scalable: they do not claim to measure fitness or experimental function directly, but they test whether the model assigns higher compatibility to natural coding sequences than to controlled perturbations.

For an autoregressive model, we define the sequence compatibility score as the length-normalized log-likelihood:

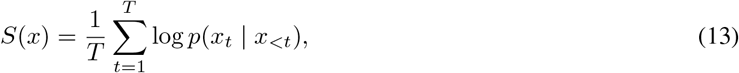

where *x*_*t*_ denotes the token at position *t*. For 6-mer models, this score can be computed either at the token level or through nucleotide-marginal likelihoods, depending on the evaluation setting.

#### 6.4.1 Triplet-Expansion Perturbation

This task is motivated by pathogenic short tandem repeat expansions, including coding CAG/polyglutamine expansions in Huntington’s disease and spinocerebellar ataxias, as well as other trinucleotide expansions such as the CAA repeat underlying Friedreich’s ataxia. Given a coding sequence *x*, we construct a perturbed sequence 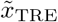 by inserting 10 consecutive CAG triplets at a controlled position within the CDS. We then compare the score of the original sequence with that of the perturbed sequence:

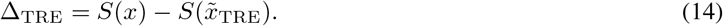

A positive margin indicates that the model assigns higher compatibility to the original CDS than to the perturbed sequence. We also report pairwise discrimination accuracy:

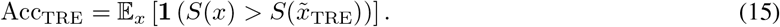

This task provides a simple and reproducible probe of whether the model recognizes that structured repeat insertion into a coding sequence can disrupt the statistical organization of natural CDS. The CAG insertion is biologically motivated by triplet-repeat expansion, but the evaluation is used here as a controlled, label-free perturbation rather than as a locus-specific disease model. Because the perturbation is easy to generate at scale, it is useful as a training-free diagnostic task for sequence-level discrimination.

#### 6.4.2 Synonymous Codon Replacement

The second perturbation task evaluates sensitivity to codon usage and natural coding-sequence statistics. Given a CDS sequence *x*, we construct a synonymous replacement 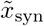 by replacing codons with synonymous alternatives. The amino acid sequence is preserved, but the nucleotide sequence distribution, codon usage pattern, local *k*-mer statistics, and species-specific coding preferences are altered. Specifically, every codon is replaced with the highest-frequency synonym for the target species, using species-specific codon usage tables from CoCoPUTs [56]. This produces a sequence that remains protein-equivalent but is less consistent with the natural DNA-level statistics of the original organism and gene context. We compute the perturbation margin

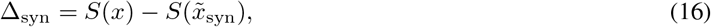

and the pairwise discrimination accuracy

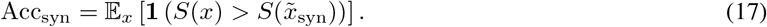

This task probes a subtler capability than triplet expansion perturbation. The synonymous sequence is not a random corruption: it preserves the encoded protein. A model must therefore detect differences in codon preference, organism-specific nucleotide composition, local coding grammar, and DNA-level constraints that are invisible at the amino acid level. At the same time, we interpret this task carefully. Synonymous mutations can be neutral, weakly selected, or functionally meaningful depending on the organism, gene, expression level, RNA structure, translation efficiency, and regulatory context. The goal is not to provide a universal measure of biological fitness, but to test whether a sequence model can distinguish natural CDS distributions from controlled protein-preserving alternatives without supervised training.

### 6.5 Long-Context Retrieval: Genomic-NIAH

#### Motivation and setup

Existing public DNA benchmarks rarely test long-range retrieval directly. Sequence Recovery and VEP probe single-position predictions; the perturbation tasks operate on a few-kbp CDS window. None of them force the model to retrieve information planted hundreds of thousands of base pairs away from the prediction position. Evo2 [9] introduced a related NIAH evaluation for DNA, but it has not been publicly released; its haystacks are constructed from random nucleotide sequences rather than real genomic context, and retrieval isn’t scored by exact-match generation. We address this with Genomic-NIAH, a publicly released needle-in-a-haystack benchmark for genomic language models, inspired by NIAH and RULER from natural-language long-context evaluation [57, 58]. Each Genomic-NIAH example plants a random (KEY, VALUE) DNA pair inside a haystack drawn from real eukaryotic genomes from OpenGenome2 [9], and asks the model to recover VALUE given the haystack followed by KEY. Because KEY and VALUE are uncorrelated with the surrounding sequence, the only way to succeed is to retrieve the planted pair at long range, turning “does the model use distal context” into a concrete generative task with a per-example accuracy. We score with exact-match retrieval.

#### Task variants

Genomic-NIAH provides four task variants of varying difficulty, inspired by RULER [58]: a plain retrieval task and three near-duplicate variants. In the near-duplicate variants, the haystack contains, in addition to the target (KEY, VALUE) pair, eight distractor pairs whose keys are obtained by applying Δ random single-base mutations to the target KEY (Δ ∈ {4, 2, 1} mutations out of 24 bp, i.e. 83%, 92%, and 96% key identity respectively); the model must retrieve the value of the exact target KEY, not any of its near-duplicates. Each task is evaluated across six context lengths (4k to 128k 6-mer tokens, equivalent to roughly 24 kbp to 786 kbp of DNA), for 24 sub-configurations in total. Each sub-configuration contains 500 examples, stratified across five needle depths within the haystack (10%, 25%, 50%, 75%, and 90% of haystack length) and four eukaryotic kingdoms.

#### Difficulty calibration across variants

Figure 5 reports Carbon performance across three of the four variants and confirms that they probe progressively finer-grained retrieval rather than the same capability at face value. Plain NIAH (no distractors) is the easiest setting, since the model only needs to retrieve the value attached to the unique KEY. The near-duplicate variant with 4 distractor mutations (83% key identity) is harder because the model must distinguish the target key from distractors that share most of its nucleotides. The 2-mutation variant (92% identity) is even harder, as the distractor keys are nearly identical to the target keys. Accuracy decreases monotonically across the three settings. We do not report the 1-mutation variant (96% key identity) in the figure; it pushes all current models to near-chance performance and is intended as a headroom benchmark for future genomic language models.

**Figure 5:**
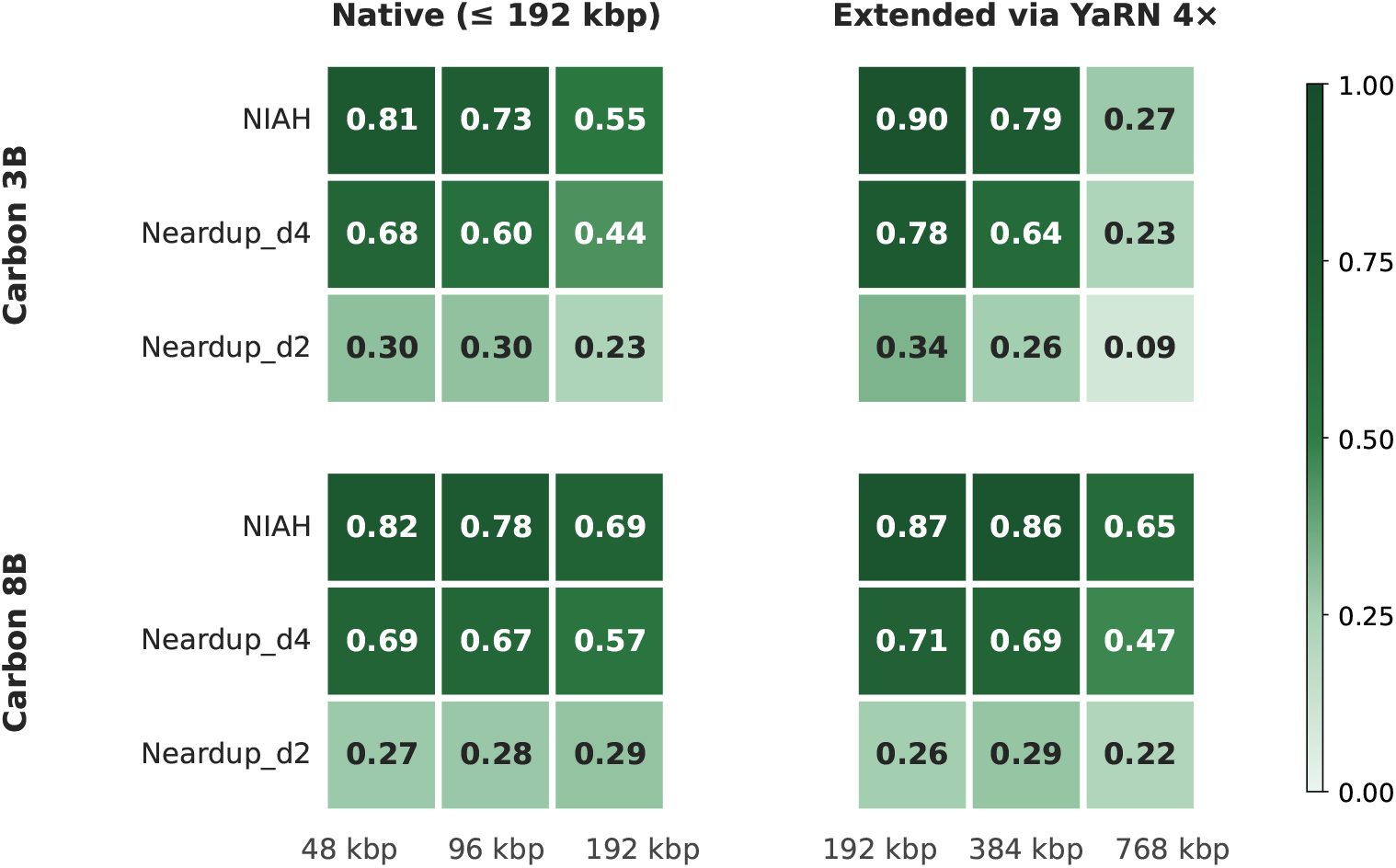
Genomic-NIAH at three difficulty levels: plain retrieval (no distractors), near-duplicates with 4 mutations in distractor keys (83% key identity), and near-duplicates with 2 mutations (92% key identity). Accuracy decreases with increasing distractor similarity, confirming that the near-duplicate variants probe progressively harder retrieval.

#### Long-context retrieval across models

Results on the plain NIAH variant are reported in Table 5 across four representative context lengths (16k to 128k tokens, 98 kbp to 786 kbp). Carbon-3B and Carbon-8B are trained natively at 32k tokens and extended at inference with YaRN to 64k (3B, 2 *×*) and 128k (8B, 4 *×*); Evo2-7B was trained natively at 1 M tokens. Evo2-7B leads at short context (16k and 32k), as expected from its much longer training context. Car-bon-3B extends cleanly with YaRN up to 64k, and Carbon-8B extends further to 128k thanks to its larger capacity; Carbon-8B is ahead of Evo2-7B at both 64k (0.86 vs 0.80) and 128k (0.65 vs 0.53). Evo2-7B is very slow at long-context inference, which forced us to evaluate it on smaller samples than Carbon. We use 500 examples per context length for Carbon, but only 100 examples at 64k and 60 at 128k for Evo2-7B; Evo2 scores were stable across runs at these reduced sample sizes.

**Table 5:** Genomic-NIAH retrieval accuracy across context lengths. Carbon columns show native context / YaRN-extended context (4*×*). Native cells are marked “–” when the context exceeds the model’s training context.

| Context length | CARBON-3B (native / YaRN) | CARBON-8B (native / YaRN) | Evo2-7B |
| --- | --- | --- | --- |
| 16 k tokens (98 kbp) | 0.73 / 0.91 | 0.78 / 0.89 | **0.97** |
| 32 k tokens (196 kbp) | 0.55 / 0.90 | 0.69 / 0.87 | **0.95** |
| 64 k tokens (393 kbp) | – / 0.79 | – / **0.86** | 0.80 |
| 128 k tokens (786 kbp) | – / 0.27 | – / **0.65** | 0.53 |

### 6.6 Efficiency Measurements

For large genomic foundation models, accuracy alone is insufficient; a model must also be practical to evaluate and deploy. For autoregressive models like Carbon, efficiency is typically measured along two main axes:

- **Throughput:** the rate at which the model can process input, typically reported in tokens (or base pairs) per second. For genomic applications this metric is particularly consequential, since downstream tasks routinely require scoring or generating sequences of 10^4^–10^6^ nucleotides, and inference cost grows quadratically with context length under standard attention.
- **Memory:** the peak GPU memory required during inference, which is dominated by model parameters and the KV cache. The KV cache scales as 𝒪(*L* · *n*_layers_ · *d*_model_) in the sequence length *L*, and at the context lengths typical of genomic modeling it frequently exceeds the parameter footprint, becoming the binding constraint on deployable batch size and maximum tractable sequence length.

These efficiency measurements are essential because tokenization choices propagate non-linearly through the cost of inference. For example, Carbon’s 6-mer tokenizer reduces the number of input tokens by a factor of six relative to single-nucleotide tokenization for the same DNA length. Because the cost of self-attention is quadratic in sequence length, this translates into roughly a 36× reduction in attention FLOPs and activation memory during prefill, and a 6× reduction in KV-cache size during autoregressive decoding. Figure 6 quantifies these gains: both Carbon-3B and Carbon-8B sustain over two orders of magnitude higher throughput than Evo2-7B at saturation, and accommodate context lengths more than an order of magnitude longer before exhausting device memory.^3^

**Figure 6:**
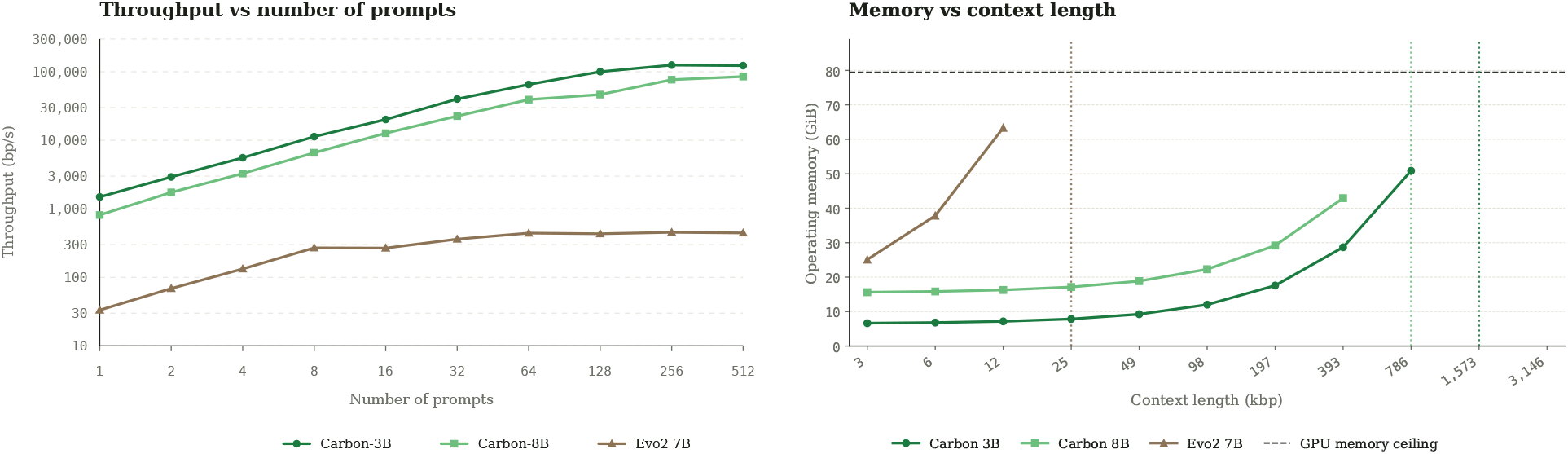
Throughput and memory efficiency of Carbon versus Evo2-7B on a single NVIDIA H100. *Left:* Throughput (base pairs per second) as a function of the number of concurrent prompts, with prefill and decode lengths held fixed at 1080 bp. *Right:* Peak operating memory (GiB) as a function of input context length at batch size 1 and a fixed decode of 30 bp; the dotted vertical lines mark the context length at which each model exhausts device memory.

### 6.7 Overall Results and Interpretation

Across this training-free evaluation suite, Carbon-3B is competitive with Evo2-7B despite having less than half the parameter count (Table 6). Carbon-3B leads on Sequence Recovery (61.54 vs 59.86, +1.68 pp) and BRCA2 (84.63 vs 83.52, +1.11 pp); the two models are essentially tied on ClinVar coding (92.89 vs 93.33) and ClinVar non-coding (91.14 vs 89.79, with Carbon-3B marginally ahead), both ClinVar splits being near saturation and noisier at end of training(Section 6.2). On Genomic-NIAH at 393 kbp, the two are also essentially tied (79 vs 80). Evo2-7B is stronger on TraitGym Mendelian and on the two controlled perturbation probes (nucleotide triplet-expansion and synonymous codon replacement). The TraitGym gap concentrates on the promoter-like (PLS) variant class; on the other TraitGym variant categories Carbon-3B is competitive with or above Evo2-7B. GENERator-v2 3B trails Carbon-3B on every reported task.

**Table 6:** Training-free evaluation results for Carbon-3B compared with GENERator-v2 3B and Evo2-7B. All scores in %. **Bold** = best per row, underlined = second best. GENERator-v2 is not reported on Genome-NIAH at 393 kbp because its maximum context is 16k tokens (96 kbp).

| Category | Task | CARBON-3B | GENERator-v2 3B | Evo2-7B |
| --- | --- | --- | --- | --- |
| Generative | Sequence Recovery eukaryote | **61.54** | 58.56 | <u>59.86</u> |
| Variant effect prediction | BRCA2 | **84.63** | 81.93 | <u>83.52</u> |
|  | TraitGym Mendelian | <u>33.65</u> | 27.91 | **37.78** |
|  | ClinVar coding (24 kb) | <u>92.89</u> | 91.55 | **93.33** |
|  | ClinVar non-coding (24 kb) | **91.14** | <u>90.13</u> | 89.79 |
| Perturbation | Nucleotide triplet-expansion | <u>85.20</u> | 83.06 | **88.43** |
|  | Synonymous codon | <u>88.89</u> | 87.03 | **91.59** |
| Long-context retrieval | Genome-NIAH @ 393 kbp | <u>79.00</u> | — | **80.00** |

Carbon-8B improves over Carbon-3B on every benchmark (Table 7). The largest gains are on Genomic-NIAH at 393 kbp (+7.0 pp), nucleotide triplet-expansion (+3.85 pp), TraitGym Mendelian (+2.78 pp), synonymous codon replacement (+2.57 pp), and Sequence Recovery (+2.51 pp); the smaller gains on ClinVar (coding +0.22, non-coding +0.49) are in part due to those tasks being near saturation at this scale.

**Table 7:** Scaling gain from Carbon-3B to Carbon-8B across all training-free benchmarks. All scores in %; Δ is the absolute gain in percentage points. **Bold** marks the larger of the two scores per row.

| Category | Task | CARBON-3B | CARBON-8B | $\Delta$ |
| --- | --- | --- | --- | --- |
| Generative | Sequence Recovery eukaryote | 61.54 | **64.05** | +2.51 |
| Variant effect prediction | BRCA2 | 84.63 | **85.72** | +1.09 |
|  | TraitGym Mendelian | 33.65 | **36.43** | +2.78 |
|  | ClinVar coding (24 kb) | 92.89 | **93.11** | +0.22 |
|  | ClinVar non-coding (24 kb) | 91.14 | **91.63** | +0.49 |
| Perturbation | Nucleotide triplet-expansion | 85.20 | **89.05** | +3.85 |
|  | Synonymous codon | 88.89 | **91.46** | +2.57 |
| Long-context retrieval | Genome-NIAH @ 393 kbp | 79.00 | **86.00** | +7.00 |

The efficiency advantage is substantial. In sequence generation and likelihood-based evaluation, Carbon achieves tens-fold faster inference than Evo2 baselines under comparable settings, while requiring several-fold less GPU memory. This advantage is crucial for practical genomic modeling. Training-free evaluation is useful only if it can be run repeatedly during model development and scaled to large benchmark sets. A model that is accurate but prohibitively expensive to evaluate is difficult to iterate on and difficult for the broader community to use.

These results reinforce the distinction between nominal context length and effective context utilization. Carbon-8B reaches 786 kbp through YaRN extension at inference, and at this context Genomic-NIAH places it ahead of Evo2-7B (0.65 vs 0.53; Table 5) despite Evo2 being natively trained at 1 M tokens. This suggests that the relevant question is not only how long a sequence a model can technically accept, but whether the model can convert its context, training data, tokenizer, and objective into measurable capability under realistic compute.

### 6.8 Scope and Future Evaluation

The eight tasks used in this report provide a practical evaluation layer across four capability axes: generation, variant effect prediction, sequence-level perturbation, and long-context retrieval. They are training-free, reproducible, scalable, and sensitive to model improvement, but they do not exhaust the biological capabilities expected from genomic foundation models. Sequence recovery and VEP probe nucleotide-level prediction; nucleotide triplet-expansion and synonymous codon replacement probe sequence-level discrimination; and Genomic-NIAH probes long-range retrieval. These tasks are useful for model development and cross-model comparison, but they are not substitutes for detailed biological validation.

Future evaluation of the Carbon series will include more biologically meaningful tasks and application-driven studies. These may include regulatory element modeling, promoter and enhancer design, splice-site prediction, geneexpression-related sequence scoring, protein-coding sequence generation, cross-species generalization, genome annotation, and experimentally validated sequence design. Such evaluations are essential for understanding how training-free capability translates into biological discovery and practical use.

These results establish that Carbon has a strong and efficient pre-trained sequence prior, that its capabilities match large baselines under controlled settings, and that its efficiency makes large-scale evaluation feasible. They do not replace more detailed biological validation; rather, they provide the foundation for it.

### 6.9 Summary

Our evaluation framework measures capability, stability, and practicality. Sequence recovery tracks intrinsic generative accuracy and serves as the primary pre-training monitor. Variant effect prediction probes nucleotide-level probabilistic reasoning. Nucleotide triplet-expansion and synonymous codon replacement provide controlled sequence-level discrimination tasks; Genomic-NIAH adds long-range retrieval on real-genome haystacks. All probes are applied without supervised fine-tuning. Efficiency measurements quantify whether the model can be evaluated and deployed at scale.

Together, these training-free tasks show that Carbon is a strong and practical generative DNA model. Carbon-3B is competitive with Evo2-7B at less than half the parameter count, while offering tens-fold inference speedups. Carbon-8B improves on Carbon-3B on every training-free task, with the largest gains on long-context retrieval, nucleotide triplet-expansion, and TraitGym Mendelian. More broadly, these results motivate a capability-driven evaluation view for genomic foundation models: progress is most informative when assessed not only by parameter count or nominal context length, but by measurable performance under efficient, reproducible, and biologically extensible evaluation.

## 7 Embedding Representation

Beyond likelihood and generation, we analyze whether Carbon learns structured internal representations of genomic sequence. A useful DNA foundation model should not only predict the next token, but also organize sequences according to biologically meaningful factors such as taxonomic background, strand orientation, coding frame, and local sequence grammar. We therefore extract hidden states from Carbon-3B and study their geometry with training-free embedding probes.

### 7.1 Data and Embedding Extraction

We compare two embeddings extracted from adjacent positions in the same input sequence. The first is the *separator embedding*, extracted from the closing DNA boundary token (`</dna>`) after the sequence. The second is the *content-token embedding*, extracted from the final DNA token immediately before the separator. Although these two hidden states differ by only one token position, they exhibit strikingly different geometries. Separator embeddings primarily capture genome-level and taxonomic structure, whereas content-token embeddings are dominated by local sequence state, including strand orientation, codon phase, and an additional major axis whose biological meaning remains unclear. The existence of such different representations at adjacent token positions suggests that autoregressive DNA models may internally organize biological information at multiple scales.

We analyze these embeddings on two held-out eukaryotic datasets. The first contains 30,000 sequences from the GenerTeam sequence-recovery task [10], covering six taxonomic groups: fungi, plants, invertebrates, protozoa, vertebrate (other), and vertebrate (mammalian). We use this dataset for separator-embedding taxonomy and context-length analysis. The second is the full held-out eukaryotic evaluation set of 29,411 sequences, covering the same six taxonomic groups. These sequences are constructed around centered coding sequence (CDS) annotations, so strand orientation and codon phase are well defined. We use this dataset for content-token embedding analysis, where the goal is to study local sequence-state variables such as strand and reading-frame structure.

We project embeddings to two dimensions with UMAP [59] and inspect the same projection under different metadata labels, including taxonomic group, strand orientation, and codon phase. Codon phase is defined as the offset between the 6-mer tokenization boundary and the canonical codon frame, with phase 0 codon-aligned and phases +1 and +2 shifted by one or two bases, respectively. For taxonomic organization, we quantify both local and global structure using K-Nearest-Neighbor (KNN [60]) accuracy, Adjusted Rand Index (ARI), and Normalized Mutual Information (NMI).

### 7.2 Separator Embeddings Capture Genome-Level Organization

We first analyze separator embeddings on balanced eukaryotic subsets at 8k, 16k, and 48k nucleotide context lengths. The goal is to test whether the model can summarize sequence context into a representation that reflects broad taxonomic identity. Importantly, the sequences in this analysis are randomly sampled held-out genomic windows rather than aligned homologous loci. They are not constructed from the same gene, orthologous region, or conserved genomic interval across species, and most pairs are unlikely to share any homologous sequence segment. Thus, taxonomic organization in this embedding space cannot be explained simply by direct sequence matching or alignment-level similarity. Instead, it suggests that the model has learned distributed genome-level regularities that recur across random genomic samples.

As shown in Table 8, all three metrics improve monotonically with context length. At 8k nucleotides, the six taxonomic groups form partially overlapping clouds; at 48k nucleotides, they resolve into more compact and distinct regions (Figure 7). This behavior is consistent with how the broad eukaryotic clades differ. Taxonomic identity is not determined by a single short motif, but by aggregate genomic properties such as GC content, codon usage, repeat density, gene architecture, splice structure, and other lineage-specific sequence statistics. Longer contexts make these distributed signals more stable, allowing the separator embedding to form a stronger sequence-level summary.

**Table 8:** Context-length ablation using separator embeddings on balanced eukaryotic subsets. Bold denotes the best result per metric.

| Context length | KNN | ARI | NMI |
| --- | --- | --- | --- |
| 8kb | 0.985 | 0.274 | 0.377 |
| 16kb | 0.990 | 0.316 | 0.415 |
| 48kb | **0.995** | **0.380** | **0.474** |

**Figure 7:**
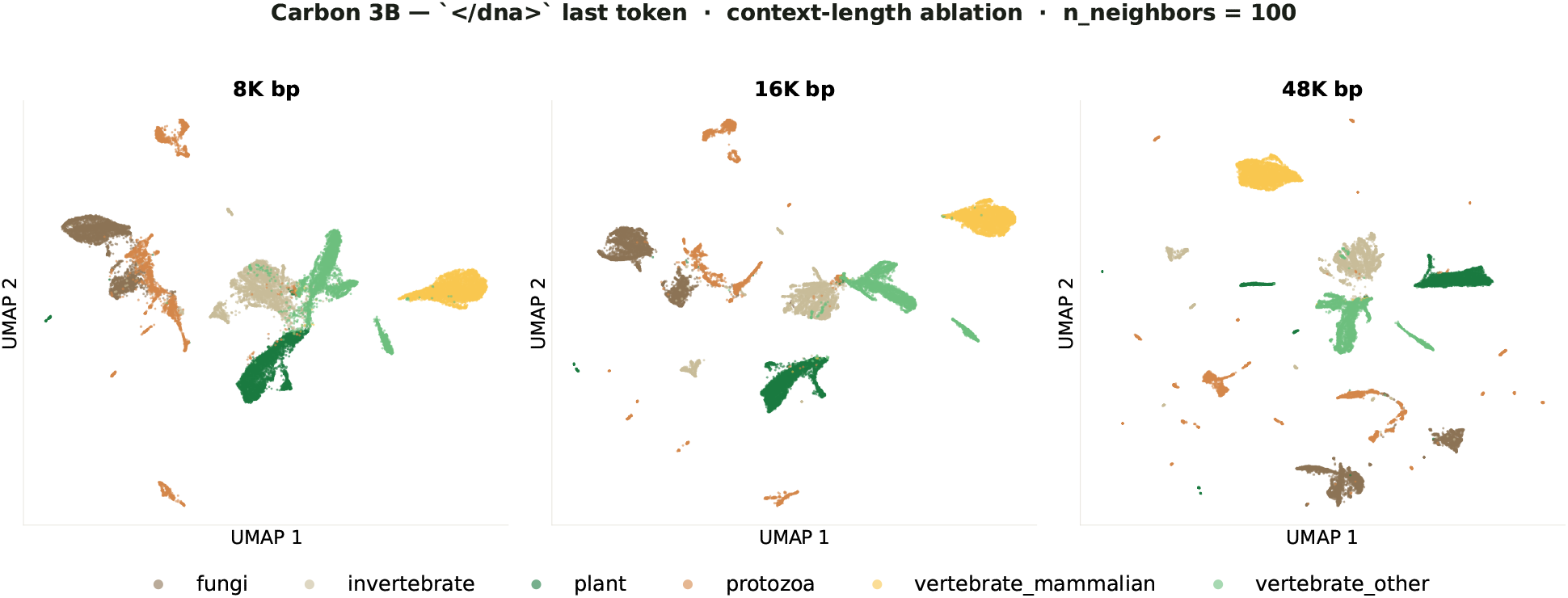
Context-length ablation. UMAP projections of separator embeddings extracted from balanced eukaryotic subsets at 8k, 16k, and 48k nucleotide context lengths, colored by taxonomic group. Each panel reports KNN, ARI, and NMI against the six-group taxonomic labels. Cluster separation improves with context length; at 48k nucleotides, mammalian vertebrates form a compact isolated region while the remaining groups become more spatially distinct.

### 7.3 Content-Token Embeddings Capture Local Sequence State

We next analyze content-token embeddings in the full 29,411-sequence eukaryotic evaluation set using a 48k nucleotide context. Each sequence is constructed so that the final DNA token falls within the centered CDS region, ensuring that the strand orientation and codon phase are well defined at the embedding position. Unlike separator embeddings, which organize strongly by broad taxonomic structure, content-token embeddings show a different geometry. When projected with UMAP, the embeddings separate into two large clusters along the first UMAP dimension.

We fit a linear SVM [61] in the (*U*_1_, *U*_2_) plane to divide the projection into two major regions and then re-embed each region independently.

The first split is not explained by strand orientation or codon phase: both labels are mixed across the two large clusters. This indicates that the dominant UMAP axis captures another source of variation. At present, we do not have a definitive biological interpretation of this axis. It may reflect a broad compositional, phylogenetic, annotation-related, or sequence-construction factor, but further controlled analyses are needed. We therefore treat this first axis as an unresolved major dimension in the content-token representation.

After separating the two large clusters with the SVM boundary, the internal structure becomes much clearer. When each group of clusters is projected independently with UMAP, strand orientation and codon phase emerge as dominant organizing factors (Figure 8). Within each major region, the embedding space separates according to forward versus reverse strand and into the three codon phases. This shows that content-token embeddings encode local coding-frame information, but that this information is not necessarily the first global axis of variation in the full embedding space.

**Figure 8:**
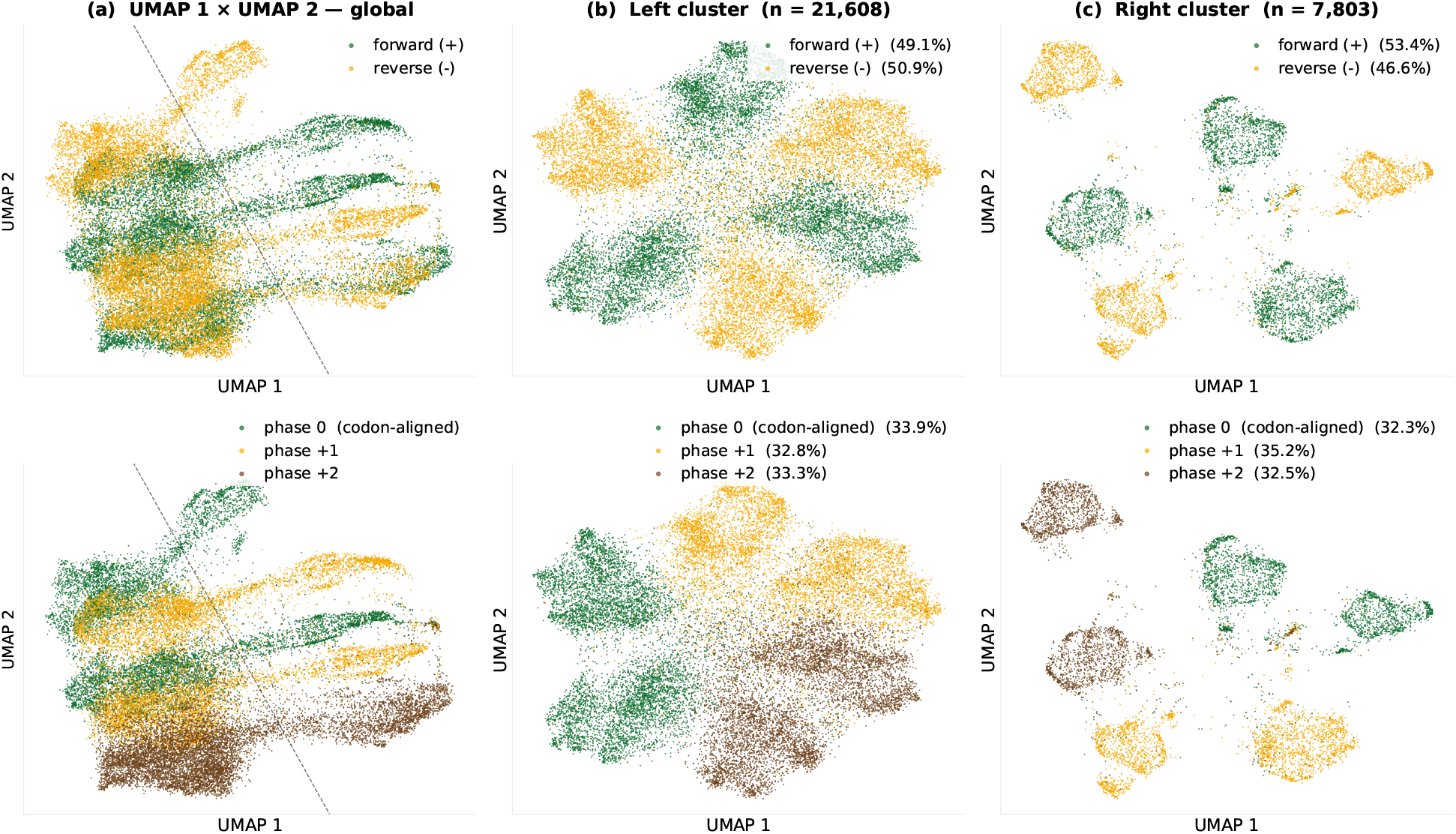
Large-scale structure of content-token embeddings. Each point is one of 29,411 held-out eukaryotic sequences. In every panel, the top row is colored by DNA strand orientation and the bottom row by codon phase. **(a)** Global view: a 2D linear SVM fit in (*U*_1_, *U*_2_) partitions the embedding into a left cluster (*n* = 21,608) and a right cluster (*n* = 7,803); the boundary is overlaid in black. Strand and codon phase are mixed across both major clusters, indicating that neither factor explains the first split. **(b)** Round-2 UMAP of the left cluster and **(c)** round-2 UMAP of the right cluster: within each cluster, strand orientation and codon phase emerge as the dominant organizing axes.

This result is notable because Carbon uses non-overlapping 6-mer tokenization. If the model were dominated by token-boundary artifacts, one might expect a 6-periodic structure. Instead, the content-token geometry recovers the natural three-phase structure of the coding sequences. This suggests that 6-mer tokenization does not prevent the model from representing nucleotide-level reading-frame information. The model learns local coding syntax at a resolution finer than that of the token unit, consistent with the role of FNS and the need for nucleotide-level reasoning.

### 7.4 Summary

These embedding analyses suggest that Carbon learns structured genomic representations rather than merely memorizing local sequence patterns. Separator embeddings organize randomly sampled non-homologous genomic windows according to broad taxonomic structure, while content-token embeddings recover biologically meaningful local variables such as strand orientation and codon phase. Both structures emerge directly from autoregressive next-token prediction pre-training, without explicit supervision for taxonomy, strand, or coding frame.

The contrast between separator and content-token embedding is particularly informative. Although the two embeddings are extracted from adjacent token positions, they encode qualitatively different biological structures. This suggests that different positions inside the autoregressive sequence can specialize according to their predictive role. Content tokens remain closely tied to local nucleotide continuation, whereas separator tokens become associated with predicting transitions between neighboring genomic segments under sequence concatenation.

This interpretation also provides a possible explanation for why separator embeddings become strongly taxonomically organized. Under concatenated training, accurately predicting the next genomic segment may require the model to infer broader contextual information such as species identity, genome organization, and lineage-specific sequence statistics. The separator token therefore becomes a natural location for aggregating genome-level information, effectively acting as an emergent summary representation despite the absence of any explicit summary objective.

More broadly, these results suggest that autoregressive DNA modeling can induce biologically meaningful organization across multiple scales simultaneously. The model learns local coding syntax, strand-aware sequence structure, and broader genome-level regularities within a unified representation space. In this sense, the embedding geometry provides representation-level evidence that Carbon is learning internal abstractions aligned with underlying biological structure, rather than simply storing surface-level sequence statistics.

## 8 Conclusion and Discussion

Carbon was developed as a practical attempt to understand what makes a generative DNA language model strong, efficient, and useful. The model itself is deliberately simple: a decoder-only autoregressive Transformer with nonoverlapping 6-mer tokenization. Most of the design effort went into the surrounding recipe: how the data was curated, how the sequence was represented, how the objective was scheduled, and how training progress was evaluated. The main observation from this work is that these choices matter substantially. A model does not need to rely only on the longest nominal context length or the largest parameter count to achieve strong performance if the training pipeline is well aligned with the statistical and biological structure of DNA.

This experience also shaped how we interpret the current stage of genomic foundation modeling. DNA modeling benefits greatly from LLM technology, but it is not simply natural language modeling with a four-letter alphabet. Genomes are noisy, redundant, sparsely constrained, unevenly annotated, and shaped by evolutionary rather than communicative pressures. In our experiments, data construction, tokenization, and objective design all produced large differences in model behavior. This suggests that the field still has substantial room for algorithmic and data-centric improvement before parameter scaling or nominal context extension becomes the dominant source of progress.

### 8.1 Dissecting the Carbon Recipe

The ablations and training runs behind Carbon suggest that its performance comes from the alignment of several design choices rather than from any single component.

The first is data. Raw genomic sequence has a different signal structure from natural language: large portions of the genome can be weakly constrained, repetitive, or only sparsely informative for the capabilities we want the model to acquire. Our data ablations show that annotation-aware mixtures outperform less curated alternatives under the same training budget, suggesting that the effective density of biological signal can matter as much as raw nucleotide count. For Carbon, data curation is therefore not merely preprocessing, but a central modeling decision.

The second is tokenization. BPE is powerful in natural language modeling and can work well in masked genomic models, but it was poorly matched to autoregressive DNA generation in our setting. DNA lacks stable word boundaries, and learned subword tokenization introduces prefix and segmentation ambiguity during next-token training. Non-overlapping 6-mer tokenization provides a simple alternative: it fixes the prediction step length, gives each continuation a unique segmentation, reduces token length by a factor of six, and improves context coverage and inference efficiency with little performance cost in our experiments.

The third is objective design. Coarse tokenization raises an immediate concern: how can a 6-mer model support nucleotide-level reasoning? FNS addresses this by marginalizing the 4,096-way 6-mer distribution into nucleotide-level probabilities, allowing the model to expose base-level supervision and perform base-level inference while retaining the efficiency of compressed tokens. At the same time, FNS is not used as a complete replacement for CE. CE is valuable early because it teaches joint 6-mer structure, while pure CE training becomes too sharp in late stages, producing the loss staircase and a widening BF16–FP32 gap in sequence recovery. The CE-to-FNS schedule uses these objectives where they are most useful: CE first learns the exact joint-token structure, and FNS later provides a smoother nucleotide-aware refinement objective that stabilizes training and improves numerical robustness.

Finally, lightweight training-free probes made this development process measurable. Sequence recovery, variant effect prediction, triplet-expansion perturbation, and synonymous codon replacement do not cover all of biology, but they are fast, reproducible, sensitive to model improvement, and applicable to frozen models without fine-tuning. In practice, they allowed us to compare tokenizers, monitor training health, detect precision issues, choose the CE-to-FNS switch point, and benchmark large baselines under controlled settings.

Taken together, the Carbon recipe is best understood as a systems-level alignment of data, representation, objective, and evaluation. Its main practical contribution is not any isolated trick, but the demonstration that a comparatively simple generative DNA model can become strong and efficient when these components are matched to the structure of genomic sequences.

### 8.2 Nominal Context and Effective Context Utilization

Long-context modeling remains one of the central challenges in genomics. Many biological mechanisms are inherently non-local: distal regulatory elements can influence gene expression across large distances, gene neighborhoods can carry functional organization, chromatin structure can couple distant loci, and evolutionary constraints may only become visible with sufficient surrounding sequence. Extending the effective context of DNA foundation models is therefore an important goal.

At the same time, our results suggest that maximum input length should not be treated as a standalone measure of model quality. A model’s nominal context length describes what it can accept, but not how much of that context contributes to prediction. Long-context capability becomes meaningful when additional sequence leads to measurable gains in sequence recovery, likelihood calibration, perturbation discrimination, representation quality, downstream performance, or long-range retrieval. This is the distinction we refer to as *nominal context length* versus *effective context utilization*.

This distinction helps interpret the Carbon results. Carbon supports a shorter maximum context than some megabase-scale models [9, 14, 62, 36], yet achieves stronger performance in our training-free evaluation suite. We do not interpret this as evidence that shorter context is inherently preferable. Rather, it suggests that model comparisons should ask how effectively a model converts available context into predictive capability under realistic compute. Efficiency is part of the same issue: genomic applications often require scoring many sequences, variants, perturbations, or genomic regions, so throughput directly affects whether a model can be used systematically. This is why we report accuracy and inference time together, and why Genomic-NIAH provides an initial step toward directly probing long-range context use.

The operational view that emerges is simple: long context is valuable when it produces measurable biological gains under practical compute constraints. The goal is not merely to advertise longer nominal inputs, but to build models that reliably transform longer genomic context into stronger biological prediction.

### 8.3 Limitations

Carbon has several limitations. First, although our evaluation suite is broader than many prior public DNA evaluations, it remains incomplete. Sequence recovery, variant effect prediction, triplet-expansion perturbation, synonymous codon replacement, and Genomic-NIAH provide useful evidence about generative quality, nucleotide-level scoring, sequence-level discrimination, training dynamics, and long-range retrieval. However, they do not cover the full range of biological applications. More realistic regulatory tasks, splicing tasks, cross-species generalization, sequence design benchmarks, and experimental validation will be needed to assess practical utility more comprehensively.

Second, annotation-aware data curation depends on available annotations. It improves signal density, but may underrepresent poorly annotated species, distal or condition-specific regulatory elements, structural variants, rare functional sequences, and biology not captured by current annotation pipelines. The current corpus is effective for Carbon, but it remains an approximation to the true functional landscape of genomes.

Third, 6-mer tokenization and FNS are practical compromises rather than final answers. Non-overlapping 6-mer tokenization improves efficiency and context coverage, but introduces phase sensitivity and represents DNA in fixed blocks. FNS recovers nucleotide-level supervision from 6-mer logits, but relies on a conditional-independence approximation among positions within each token. This approximation improves late-stage stability, but is not always biologically correct, especially in coding regions, splice sites, and regulatory motifs where nucleotide combinations can be functionally critical. This explains why FNS from scratch is weaker than CE-to-FNS switching in our experiments.

Finally, Carbon remains primarily a DNA sequence model. It can generate sequences, score likelihoods, and support training-free evaluations, but it does not yet provide a natural interface for people to ask biological questions, inspect model reasoning, or translate sequence-level predictions into mechanistic explanations. This modality gap limits interpretability, controllability, and accessibility, and motivates the next stage of the Carbon project.

### 8.4 Future Directions

The next stage of Carbon will move in two connected directions. The first is to continue improving DNA sequence modeling itself. This includes better data mixtures, stronger annotation-aware and annotation-free curation, improved tokenization, more stable objectives, better context utilization, and broader training-free and downstream evaluations. We are especially interested in measuring long-context benefit directly: not only whether a model accepts longer inputs, but whether additional context improves biological prediction under realistic compute.

The second direction is deeper integration between DNA language and natural language. A more useful genomic foundation model should allow users to describe biological goals in natural language and receive sequence-level predictions, explanations, and experimentally testable designs. Users may want to design a genetic circuit for context-specific gene expression, tune a promoter, generate a sequence implementing a desired function, interpret a deleterious variant, assess a synonymous mutation, or identify the sequence features supporting a regulatory prediction. Such tasks require more than accurate sequence likelihoods: they require connecting genomic sequence modeling with natural-language reasoning, biological annotations, functional assays, evolutionary information, and domain knowledge.

This direction also motivates a broader view of how future models can be evaluated. Beyond benchmark metrics, the value of a genomic foundation model will increasingly depend on whether it helps people solve meaningful biological problems and generate experimentally testable hypotheses. Important application-oriented settings include regulatory element design, promoter and enhancer engineering, splice-aware modeling, variant interpretation in non-model organisms, protein-coding DNA generation, genome annotation, and organism-specific sequence optimization.

If realized, this integration could become a transformative interface for biological understanding and engineering, making sequence-level reasoning, mechanistic interpretation, and biological design more programmable, customizable, and accessible. Instead of reducing each biological problem to a specialized model, scoring function, or manually engineered pipeline, users could express goals, mechanisms, constraints, and experimental feedback directly in natural language, while the model translates them into interpretable hypotheses and experimentally testable DNA designs.

Carbon marks a milestone for efficient generative DNA modeling, but it is still an early step. The main result is not that a particular 6-mer Transformer is the final architecture for DNA. Rather, the Carbon experience suggests that careful alignment between modeling choices and the biological structure of DNA can substantially change the performance–efficiency frontier. Future Carbon models will continue exploring how genomic sequence modeling can become more capable, interpretable, and usable across scientific and applied settings.

## A Training Objective

This appendix expands the objective-level discussion in Section 5 and provides the technical details omitted from the main text. We first review exact 6-mer cross-entropy and the loss staircase observed during CE-only training. We then introduce Factorized Nucleotide Supervision (FNS), describe the CE-to-FNS objective schedule, and discuss its relation to base-pair-level inference, label smoothing, multi-token prediction, and blockwise attention.

### A.1 Exact 6-mer Cross-Entropy

Let ℬ = {*A, C, G, T*} denote the nucleotide alphabet and let 𝒱 = ℬ^6^ be the 6-mer vocabulary, with |𝒱| = 4^6^ = 4096. At autoregressive step *t*, the model receives previous tokens as context and produces logits *z*_*t*_ ∈ ℝ^|𝒱|^, which define a probability distribution over 6-mer tokens:

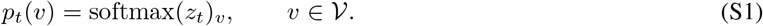

Given the ground-truth 6-mer token *y*_*t*_ ∈ 𝒱, standard next-token cross-entropy is

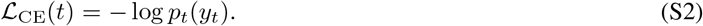

This objective treats each 6-mer as an atomic class. Only the exact target token contributes to the positive learning signal; all other tokens are treated as incorrect. For example, if the target token is `TATATA`, then `TATATT`, which matches five out of six nucleotides, receives no more credit than a completely different token such as `CGCGCG`. The gradient of cross-entropy with respect to the logit of token *v* is

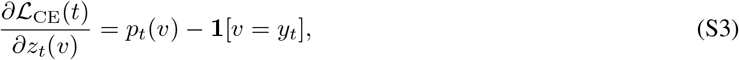

so every non-target token is pushed down regardless of its nucleotide-level similarity to the target.

This all-or-nothing supervision can be too strict for DNA. Functional DNA elements are not always defined by exact strings. Promoter motifs, splice signals, transcription factor binding sites, and other regulatory elements often tolerate substitutions, insertions, deletions, phase shifts, and context-dependent variation. The TATA box provides a simple example. Although its name suggests a canonical pattern such as `TATATA`, functional TATA-like elements form a degenerate family rather than a single fixed sequence. A nearby variant such as `TATATT` may still be compatible with TATA-box function in the appropriate sequence context, but exact 6-mer CE treats it no differently from a completely unrelated token. In such regions, forcing the model to concentrate all probability mass on one exact 6-mer can overpenalize biologically plausible near variants.

At the same time, cross-entropy is not simply wrong. It is valuable because it trains the joint distribution of nucleotides within a 6-mer. This is important for coding regions, splice sites, codon structure, and other contexts where nucleotide combinations matter. Thus, the issue is not that CE should be discarded, but that exact-token CE can become too sharp as the only objective throughout the entire training process.

### A.2 The Loss Staircase Phenomenon

During Carbon pre-training, we observed a recurring instability under pure CE training. The training loss does not merely exhibit short transient spikes. Instead, the smoothed loss can suddenly increase and then remain at a higher level for an extended period. We refer to this behavior as a *loss staircase*. Unlike ordinary optimization spikes, a loss staircase represents a persistent upward shift in the training trajectory. Each staircase is preceded by a steady rise in gradient norm over many steps, suggesting the model is already drifting into a fragile regime before the loss visibly shifts. We tried several standard mitigations: resuming training from an earlier batch, tightening gradient clipping, and lowering the learning rate. None prevented the staircase outright; some only delayed when it appeared. This pattern argues against a data-side cause. Figure S1 shows the training-loss and gradient-norm trajectories for Carbon-3B and Carbon-8B under pure CE training in preliminary runs.

We hypothesize that this phenomenon reflects a late-stage mismatch between exact 6-mer supervision and the intrinsic ambiguity of genomic sequence. In early training, CE provides a strong and useful signal: the model learns local sequence grammar, codon structure, motif composition, species-specific sequence patterns, and dependencies among nucleotides within each 6-mer. However, after these coarse and intermediate structures have been learned, continued CE optimization increasingly forces the model to resolve exact 6-mer identities in contexts where multiple near variants may be plausible. Across similar contexts, the model may receive conflicting exact-match gradients, especially in noisy, redundant, or weakly constrained genomic regions. This can push the model into a brittle regime where further optimization becomes unstable.

**Figure S1:**
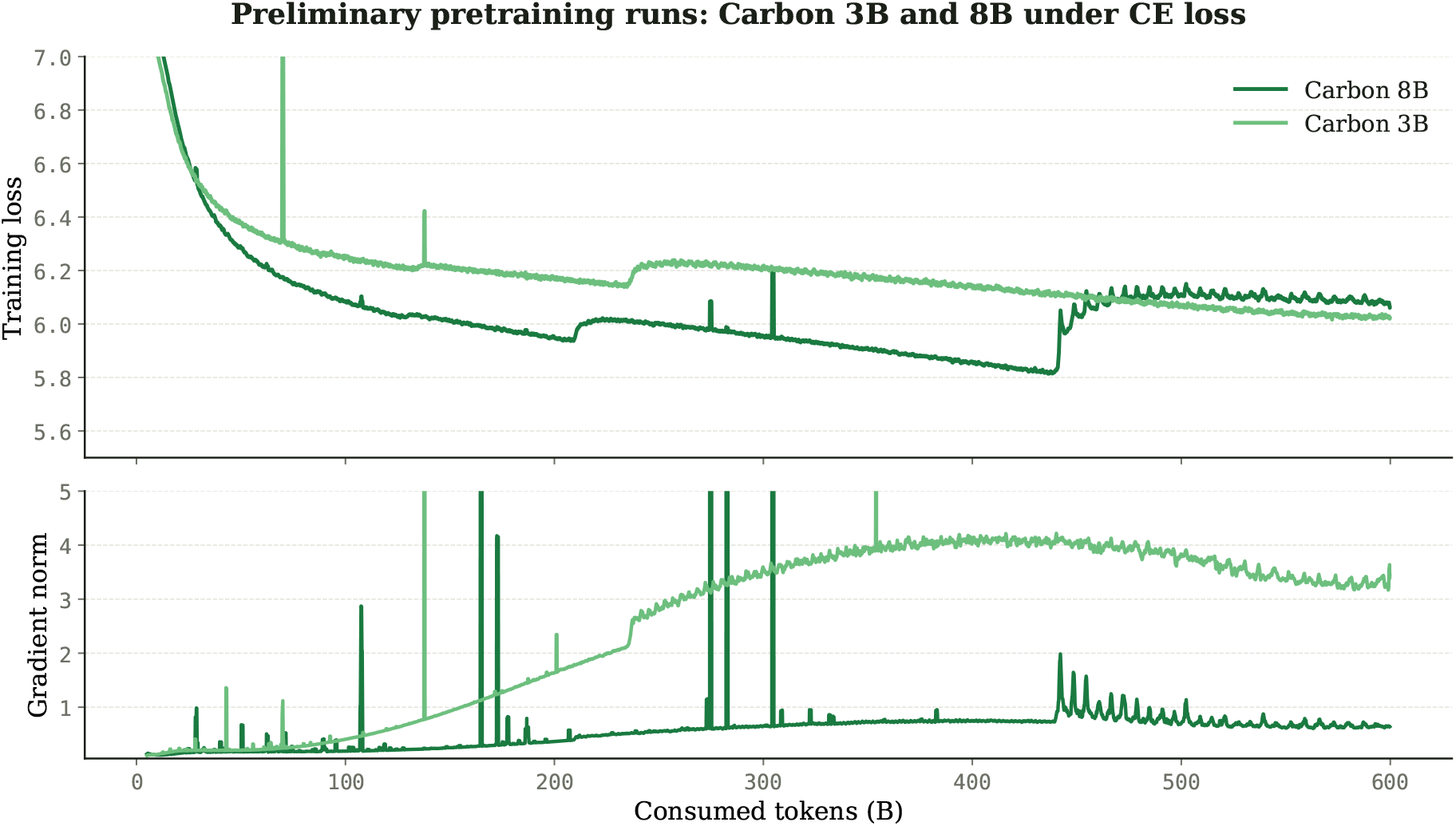
Training dynamics under pure CE training for Carbon-3B and Carbon-8B during preliminary 600 B tokens of pretraining. The smoothed training loss shows the loss staircase behavior, which is especially pronounced in the 8B model: at the jump, the 8B CE loss returns to roughly 3B level despite the larger capacity. Gradient norms rise steadily in the steps preceding each staircase, suggesting the model drifts into a fragile regime before the loss visibly shifts.

This interpretation remains a hypothesis. A complete explanation would require more detailed analyses of token entropy, gradient conflict, genomic region type, and local sequence degeneracy. Nevertheless, the empirical signal is clear: pure CE training can enter a late-stage regime in which optimization becomes less stable, and model behavior becomes more sensitive to numerical precision.

### A.3 Tracking Training Dynamics with Training-Free Probes

We evaluate intermediate checkpoints throughout pre-training using lightweight training probes. These probes are used alongside training loss to track capability growth, objective behavior, and numerical robustness as optimization progresses. The full probe suite, including sequence recovery, variant effect prediction, triplet-expansion perturbation, and synonymous codon replacement, is described in Section 6. Here, we focus on sequence recovery because it is the most sensitive diagnostic for the CE-related dynamics discussed above, especially the loss staircase and the BF16/FP32 gap.

Raw training loss alone is insufficient for this purpose. Loss values are tokenizer-dependent and can be affected by repetitive, low-complexity, or weakly constrained regions in the training corpus. A checkpoint may therefore show favorable loss trends without necessarily improving in generative capability. Training-free probes provide a complementary signal: they are inexpensive to run, require no fine-tuning, and evaluate frozen checkpoints under a fixed inference protocol.

Sequence recovery (described in full in Section 6) is well-suited for this purpose. Given a fixed-length DNA prompt, the model generates a fixed number of subsequent nucleotides, and the continuation is compared with the ground-truth sequence at nucleotide resolution. Because both prompt and generation length are defined in base pairs rather than tokenizer-specific units, sequence recovery provides a tokenizer-agnostic measure of intrinsic generative quality and is sensitive to degradation when optimization becomes unstable. It is genuinely generative (the model has to produce the continuation, not just provide representations for a classifier) and remains non-saturating over a long training horizon. We therefore use it as the primary monitoring signal in this section, analogous in spirit to perplexity tracking in natural-language modeling but more comparable across tokenization schemes.

Most importantly for the objective analysis, sequence recovery revealed a numerical signature of the loss staircase. Before the staircase, BF16 and FP32 inference produce closely aligned recovery accuracy. After the staircase, however, BF16 performance can drop substantially, while FP32 performance may remain stable or even continue to improve. This indicates that pure CE training can push the model into a regime where correct generation depends on fine probability distinctions among many 6-mer classes. Such distinctions may still be recoverable under FP32, but become fragile under BF16 inference.

This precision sensitivity has direct practical consequences. A model that requires FP32 inference to maintain its generative capability is less useful for large-scale genomic applications, because FP32 increases memory consumption and reduces throughput. One of the main motivations for 6-mer tokenization is efficient inference; losing this advantage through precision fragility would weaken the practical value of the model. We therefore treat the BF16/FP32 gap in sequence recovery as an important signal of training dynamics rather than a minor implementation detail.

### A.4 Factorized Nucleotide Supervision

Motivated by the late-stage CE instability and the BF16/FP32 gap, we replace the all-or-nothing exact-token objective with Factorized Nucleotide Supervision (FNS) [17]. The key idea is to preserve the 6-mer output space while exposing nucleotide-level supervision through probability marginalization. The model still produces logits over the 4,096 6-mer tokens, but the loss is computed from the nucleotide marginals induced by this distribution.

For each nucleotide position *j* ∈ {1, &, 6} and base *b* ∈ ℬ, we define the marginal probability

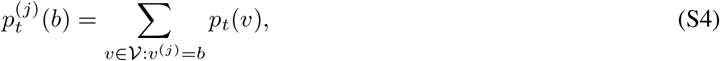

where *v*^(*j*)^ denotes the *j*-th nucleotide of token *v*. Given the target token

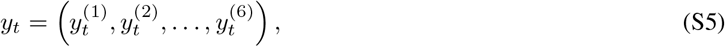

the FNS loss is

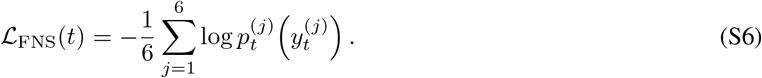

For simplicity, the FNS loss is applied uniformly to all six positions of every 6-mer token, including the rare tail 6-mer at a sequence boundary whose final positions may correspond to padding rather than valid bases; this approximation affects at most one token per sequence and has a negligible effect on the average loss.

FNS introduces structured partial credit. A token that matches five out of six nucleotides contributes probability mass to five correct nucleotide marginals, whereas a completely mismatched token contributes little or none. The gradient of FNS with respect to a token logit is

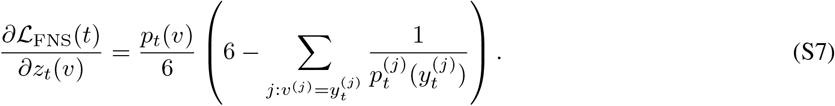

Unlike CE, this gradient depends on how many nucleotide positions in *v* match the target. Near-miss tokens are therefore not treated identically to completely incorrect tokens, giving the model a smoother optimization landscape and a denser learning signal.

FNS can also be interpreted as a blockwise nucleotide predictor. Given the hidden state *h*_*t*_, the 6-mer distribution induces six nucleotide-level categorical distributions:

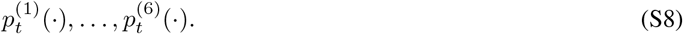

Under a conditional-independence approximation among the six nucleotide positions given *h*_*t*_, these marginals define

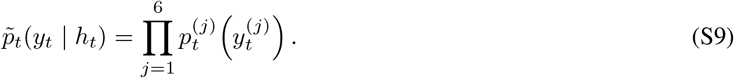

The FNS objective is then the normalized negative log-likelihood of this blockwise single-nucleotide predictor:

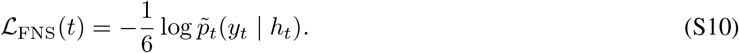

Conceptually, FNS makes a 6-mer model predict six nucleotide targets in parallel from a shared hidden state, while preserving the computational efficiency of 6-mer tokenization.

### A.5 FNS as a Late-Stage Refinement Objective

Although FNS stabilizes training, using FNS from the beginning is not optimal in our experiments. FNS-from-scratch removes the loss staircase, but the final model underperforms a CE-trained model at the same training budget. This suggests that the conditional-independence approximation in FNS is useful for stability but incomplete as a full training objective.

The reason is that nucleotides within a 6-mer are not independent in many biologically important contexts. In coding regions, a single nucleotide substitution can change an amino acid, create a premature stop codon, or disrupt codon-level constraints. In splice sites, promoter motifs, and regulatory elements, combinations of nucleotides may matter more than individual positions. In these cases, giving partial credit to a near-miss token can be biologically inappropriate: a 5-out-of-6 match may still represent a functionally severe error.

CE is therefore important in early training because it teaches the joint 6-mer distribution. It encourages the model to learn dependencies among nucleotide positions inside each token. FNS, by contrast, decomposes the target into nucleotide marginals and can underemphasize within-token dependencies if applied too early. This creates a natural division of labor: CE is useful for learning sharp joint structure, while FNS is useful for smoothing late-stage optimization and improving nucleotide-level calibration.

This relationship can be expressed in information-theoretic terms. CE trains the model toward the joint target distribution *P* (*Y* ^(1)^, &, *Y* ^(6)^|*h*), whereas FNS trains the six marginal distributions *P* (*Y* ^(*j*)^|*h*). The joint entropy satisfies

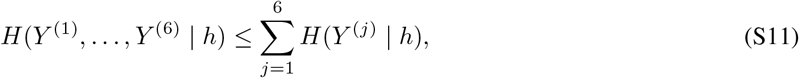

with equality only when the six positions are conditionally independent. The gap corresponds to conditional mutual information among positions within the 6-mer. Early in training, learning this dependency structure is important. Late in training, once much of this structure has been learned, marginal supervision can provide a smoother and more stable refinement signal.

### A.6 CE-to-FNS Objective Switching

Based on these observations, Carbon adopts a staged objective schedule. The model is first trained with standard CE, allowing it to learn joint 6-mer structure and conditional dependencies. After the model approaches the regime where CE loss staircases begin to appear, we reduce the learning rate and switch the objective to FNS. The second stage continues training with smoother nucleotide-level supervision.

Let *T*_switch_ denote the switching step. The training objective is

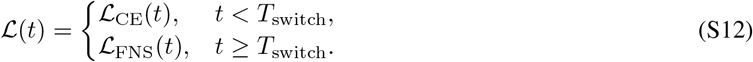

At the switch point, we also reduce the learning rate:

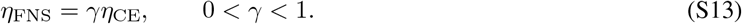

Concrete values for *T*_switch_ and *γ* used in Carbon-3B and Carbon-8B are given in the pretraining recipe below.

The intuition is that CE first teaches the model a strong joint-token prior. Once this prior has formed, FNS relaxes the overly strict exact-token penalty and prevents the model from being forced into brittle distinctions among near-equivalent 6-mers. Because the model has already learned useful conditional dependencies during the CE stage, the conditional-independence approximation of FNS is less damaging in the second stage than it would be from scratch.

FNS does not change the location of the ideal optimum for a deterministic target. The FNS loss is minimized when

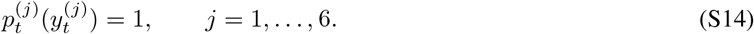

The only 6-mer token that satisfies all six nucleotide constraints simultaneously is the exact target token *y*_*t*_. Therefore, assigning all probability mass to *y*_*t*_ remains the optimum. FNS does not make incorrect tokens equally desirable; it changes the optimization path by providing structured partial credit before the exact solution is reached.

This is why CE-to-FNS switching works better than FNS-from-scratch. The CE stage learns joint 6-mer dependencies; the FNS stage then smooths the late-stage objective without discarding the learned structure, because the exact target token still receives the maximum reward. In this sense, the switch acts as an objective curriculum: hard joint-token learning first, nucleotide-aware refinement second.

### A.7 Switch Timing and Precision Robustness

The timing of the CE-to-FNS switch is empirically important. Switching too early resembles FNS-from-scratch: the model receives smoother gradients but does not learn joint 6-mer dependencies as effectively, leading to weaker final performance. Switching too late reduces the instability but may not fully remove the BF16/FP32 gap, suggesting that some precision-sensitive probability geometry has already been learned. The best results are obtained when the switch occurs shortly before the loss staircase, or at the earliest signs of unstable CE dynamics.

This timing dependence is closely related to numerical precision. Under CE, the model solves a 4,096-way classification problem at every step. Even when the correct token is among the top candidates, its probability may remain relatively small because probability mass is distributed across many plausible 6-mer continuations. Decoding and ranking can therefore depend on fine distinctions among small token probabilities. These distinctions are sensitive to reduced-precision inference.

FNS transforms the same 4,096-way distribution into six 4-way nucleotide distributions. Each nucleotide marginal operates at a larger probability scale. Instead of distinguishing many small token probabilities, the model is trained to assign probability among four bases at each position. This makes the objective more tolerant to numerical precision and helps align BF16 inference with FP32 inference. Empirically, after switching to FNS at the appropriate time, BF16 sequence recovery closely matches the corresponding FP32 performance, while continued CE training retains a larger precision gap after the loss staircase.

This observation provides a practical reason for using FNS beyond its biological interpretation. FNS not only introduces nucleotide-level supervision but also improves the numerical robustness of 6-mer training and inference. For a model designed to be efficient at scale, this matters: BF16 robustness directly affects memory usage, throughput, and the feasibility of large-scale training-free evaluation.

### A.8 Relation to Label Smoothing, Multi-Token Prediction, and Blockwise Attention

FNS is related to, but distinct from, label smoothing [63]. Standard label smoothing assigns some probability mass to non-target classes, typically uniformly or according to a heuristic distribution. It does not use the internal structure of the label space. FNS, by contrast, defines partial credit through nucleotide identity. A token receives support only insofar as it shares nucleotides with the target at the corresponding positions. The smoothing is therefore structured, deterministic, and biologically interpretable.

FNS is also, in spirit, related to multi-token prediction [64]. A 6-mer model can be viewed as reading and predicting DNA in blocks of six nucleotides. Under CE, each block is treated as a single atomic token. Under FNS, the same block is factorized into six nucleotide-level predictions from a shared hidden state. Thus, FNS provides a form of blockwise multi-nucleotide supervision that preserves the efficiency of compressed tokenization while recovering part of the resolution of single-nucleotide modeling.

However, FNS does not make a 6-mer Transformer equivalent to a single-nucleotide Transformer. There are two main differences. First, FNS relies on a conditional-independence approximation among the six nucleotide positions within a token, whereas a single-nucleotide autoregressive model predicts nucleotides sequentially and can model these dependencies directly through the autoregressive factorization. Second, the hidden state used by FNS is computed at the 6-mer block level. All six nucleotide marginals predicted by FNS share the same token-level context representation and the same attention pattern over previous 6-mer blocks. By contrast, a single-nucleotide Transformer has a separate hidden state and attention pattern for each nucleotide position. In this sense, 6-mer tokenization with FNS is not only factorized within each block, but also attends to previous sequence at block granularity [65].

This distinction clarifies the trade-off. A Transformer with single-nucleotide tokenization and CE is mechanistically cleaner: it avoids the conditional-independence approximation and provides full nucleotide-level attention resolution. However, it is computationally expensive because the sequence length is six times larger for the same DNA span, and the attention cost during prefill can be up to 36 times larger. A 6-mer Transformer with FNS is therefore not a perfect substitute for single-nucleotide modeling, but a practical and efficient approximation. It sacrifices some within-block and attention-granularity expressiveness in exchange for a large gain in context coverage, memory efficiency, and inference speed. In Carbon, this trade-off proved favorable empirically.

### A.9 FNS for Base-Pair-Level Inference

The same probability marginalization used in FNS training can also be used at inference time. This gives a 6-mer model two base-pair-level inference modes: base-pair-level generation and base-pair-level sequence scoring. Both modes use the original 4,096-way 6-mer distribution, but interpret it through nucleotide marginals rather than through the probability of a single atomic 6-mer token.

#### Base-pair-level generation

In direct token-level generation, the model selects or samples a full 6-mer token from the distribution *p*_*t*_(*v*). This means that the next six bases are determined by one coarse token decision. In base-pair-level generation, we instead marginalize the 6-mer distribution into six nucleotide distributions:

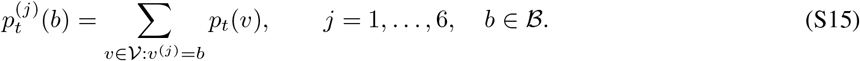

Under greedy decoding, we choose the most likely base at each position,

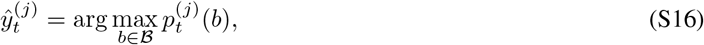

and then recombine the six selected bases into the corresponding 6-mer token 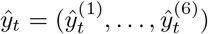. Under sampling, each base is sampled independently from its corresponding marginal distribution before recombination. The resulting 6-mer token is then appended to the context, and autoregressive generation proceeds as usual.

This procedure is different from simply choosing the most likely 6-mer token. A token with the highest joint 6-mer probability is not necessarily the token formed by the highest-probability base at each of the six positions. Base-pair-level generation therefore follows the factorized nucleotide view used by FNS: it asks the model to make six nucleotide decisions from the same hidden state, rather than one atomic 6-mer decision.

#### Base-pair-level sequence scoring

The same marginalization also defines a nucleotide-resolved likelihood for a given DNA sequence. For a target 6-mer token

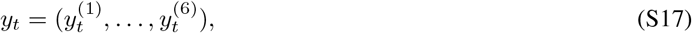

the base-pair-level score is computed from the nucleotide marginals:

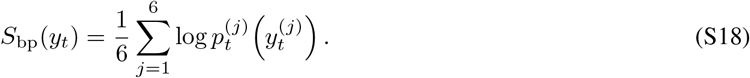

For a full sequence, we average this quantity over all valid nucleotide positions:

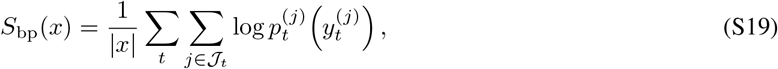

where 𝒥_*t*_ denotes the valid nucleotide positions within token *t* and excludes padding positions at sequence boundaries when needed. This produces a probability score for every base in the sequence, even though the model itself operates over 6-mer tokens.

Base-pair-level scoring is especially important for variant effect prediction and perturbation analysis. For a singlenucleotide variant, the reference and alternate alleles can be scored directly by their marginal probabilities at the affected within-token position. For sequence-level perturbation tasks, the score of the reference or perturbed sequence can be computed as the average nucleotide-marginal log-likelihood. This is the inference-time mechanism that reconciles coarse 6-mer tokenization with single-nucleotide resolution.

#### Empirical effect

Table S1 compares direct token-level inference with FNS-based base-pair-level inference. Across sequence recovery, variant effect prediction, and perturbation-based evaluations, base-pair-level inference improves performance over direct token-level generation or scoring. This supports the view that the nucleotide marginals learned through FNS are not only useful for training stability, but also provide a better inference interface for DNA tasks that are naturally defined at base-pair resolution. The main cost is computational: marginalizing the 4,096-way distribution into six nucleotide distributions introduces approximately 35% additional inference time in our implementation. This gives users a flexible choice between direct 6-mer inference, which maximizes speed, and FNS-based base-pair-level inference, which provides stronger nucleotide-resolution generation and scoring for tasks where base-level probabilities are important.

## B Ablation Curves

This appendix section collects the intermediate-checkpoint curves for the data and tokenizer ablations referenced in the main body. Each panel is a 2 × 2 grid of training-free metrics evaluated every ~8.3 B tokens (24 evaluation points over a 50 B-token run): Sequence Recovery (SR), BRCA2 AUROC, nucleotide triplet-expansion discrimination, and synonymous-codon discrimination. ClinVar and TraitGym Mendelian are held out from the ablation-selection signal and reported only on final models in Section 6, to avoid overfitting ablation choices to those specific evaluation splits. Numbers cited in figure captions are at step-24 k (50 B tokens) unless otherwise noted.

**Table S1:** Effect of FNS nucleotide-marginal inference on training-free benchmarks. **token**: standard 6-mer autoregressive decoding / token-level likelihood scoring. **bp**: FNS-marginalised base-pair-level generation and scoring. Δ = bp −token. Task scores in %; throughput in kbp/s (single A100-80 GB, vLLM [66], dynamic batching, *n*=16 prompts, 1 kbp output). Throughput Δ is the bp/token ratio (1.0 = no overhead). **Bold** = best per row, underlined = second best.

| Category | Task | CARBON-3B |  |  | CARBON-8B |  |  |
| --- | --- | --- | --- | --- | --- | --- | --- |
| | | token | bp | $\Delta$ | token | bp | $\Delta$ |
| Generative | Sequence Recovery eukaryote | 61.5 | 62.4 | +0.9 | <u>64.0</u> | **64.9** | +0.9 |
| Variant effect | BRCA2 | 84.6 | 84.6 | 0.0 | 85.7 | **85.8** | +0.1 |
|  | TraitGym Mendelian | 33.7 | <u>34.5</u> | +0.8 | **36.4** | 33.8 | −2.6 |
| Perturbation | Nucleotide triplet-expansion | 85.2 | 87.2 | +2.0 | <u>89.0</u> | **90.8** | +1.8 |
|  | Synonymous codon | 88.8 | 90.2 | +1.3 | <u>91.4</u> | **92.8** | +1.4 |
| Throughput | kbp/s | 18.9 | 12.3 | 0.65× | 12.0 | 12.3 | 1.0× |

### B.1 Pre-training Data Source: GENERator vs OpenGenome2 eukaryote subset

**Figure S2:**
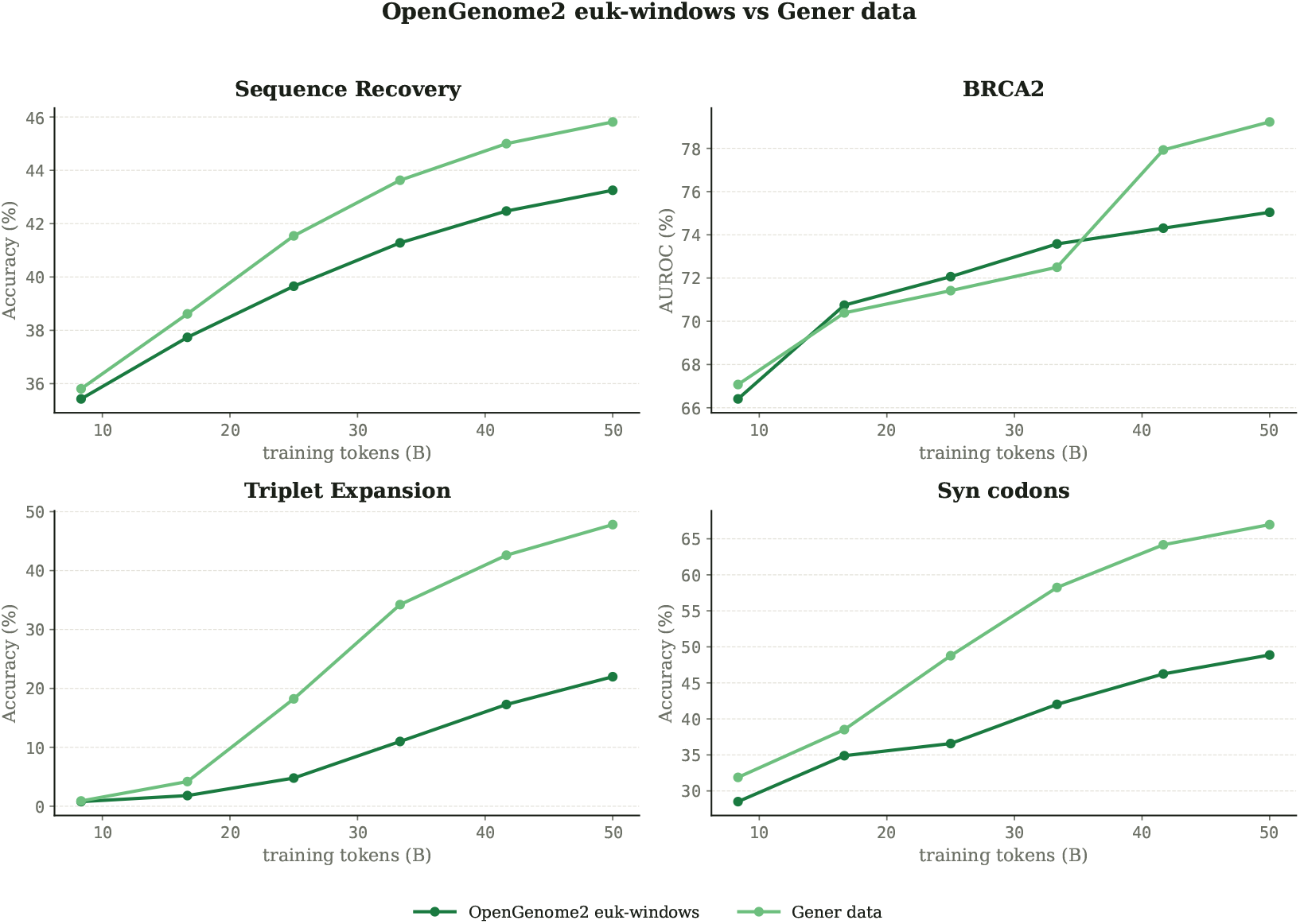
Pre-training data source under matched 6-mer / 50 B-token compute. Gener data beats the OpenGenome2 eukaryote subset on every metric, with the largest gaps on perturbation tasks (Triplet Expansion +26 pp, Synonymous codon +18 pp).

### B.2 Sequence Construction: raw vs concatenated windows

**Figure S3:**
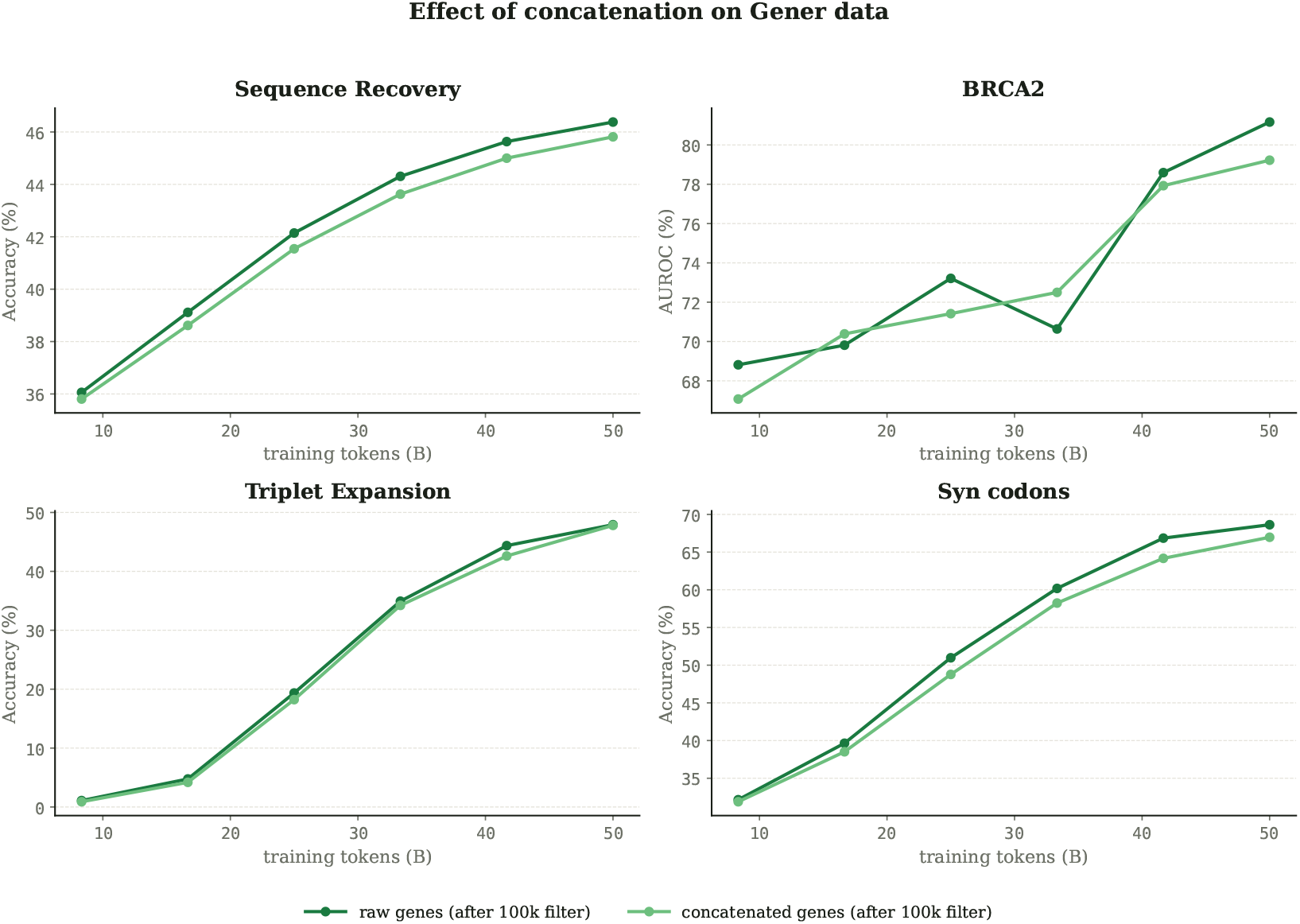
Raw functional windows vs concatenated long windows at the 8 k base-pretraining context (GENERator, 100 kbp length filter, 6-mer). Raw windows are slightly better on every metric (Δ ≈ 0.6–2 pp). Concatenation is useful when long training sequences are needed, so we defer it to the long-context extension phase (Section 5).

### B.3 Length Filter

The Gener corpus contains a long tail of very long contigs, much of which is weakly constrained background sequence (Section 3). We test three length cutoffs at matched compute: unfiltered, 200kbp, and 100 kbp. Figure S4 reports the training-free curves across the four ablation metrics.

### B.4 mRNA and mRNA-splice Mixes

Transcript-level data from OpenGenome2 (mRNA and mRNA-splice) provides coverage of splice-junction context that is less present in the Gener backbone. We compare the two subsets at a 20% mix share against a Gener-only baseline at matched compute. Figure S5 shows the per-task curves over training.

## C FNS-phase Training Stability

Here we show the training loss trajectory for both Carbon-3B and Carbon-8B after the CE-to-FNS switch at 100 B tokens through the end of pretraining, supporting the claim in Section 5 that the switch removes the late-stage loss staircase observed under prolonged CE training.

**Figure S4:**
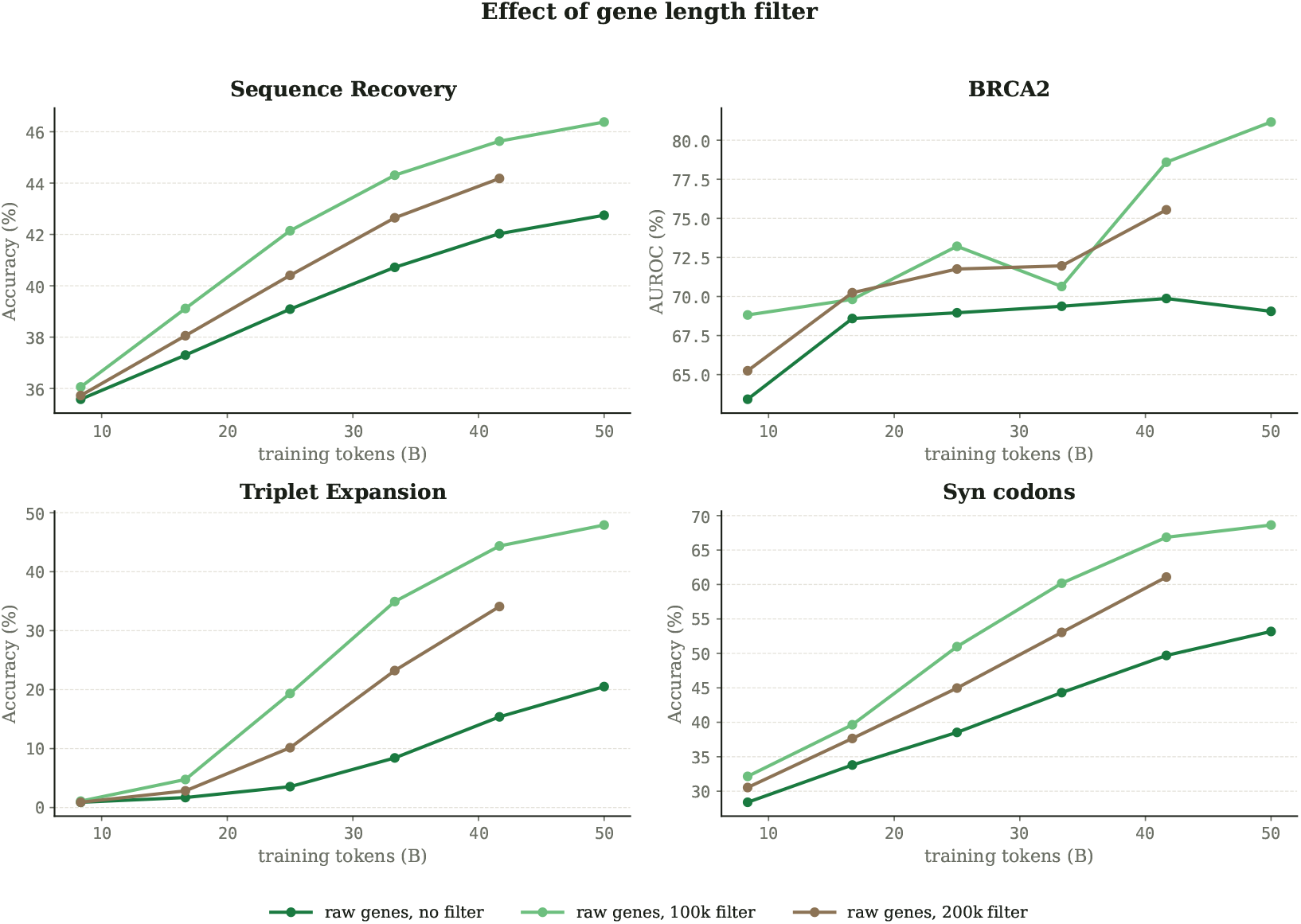
Length filter on the GENERator corpus (no concatenation, 6-mer). Dropping sequences longer than 100 kbp wins on every metric: the unfiltered corpus is dominated by very long, mostly background regions that dilute the training signal, and 200 kbp sits between unfiltered and 100 kbp.

**Figure S5:**
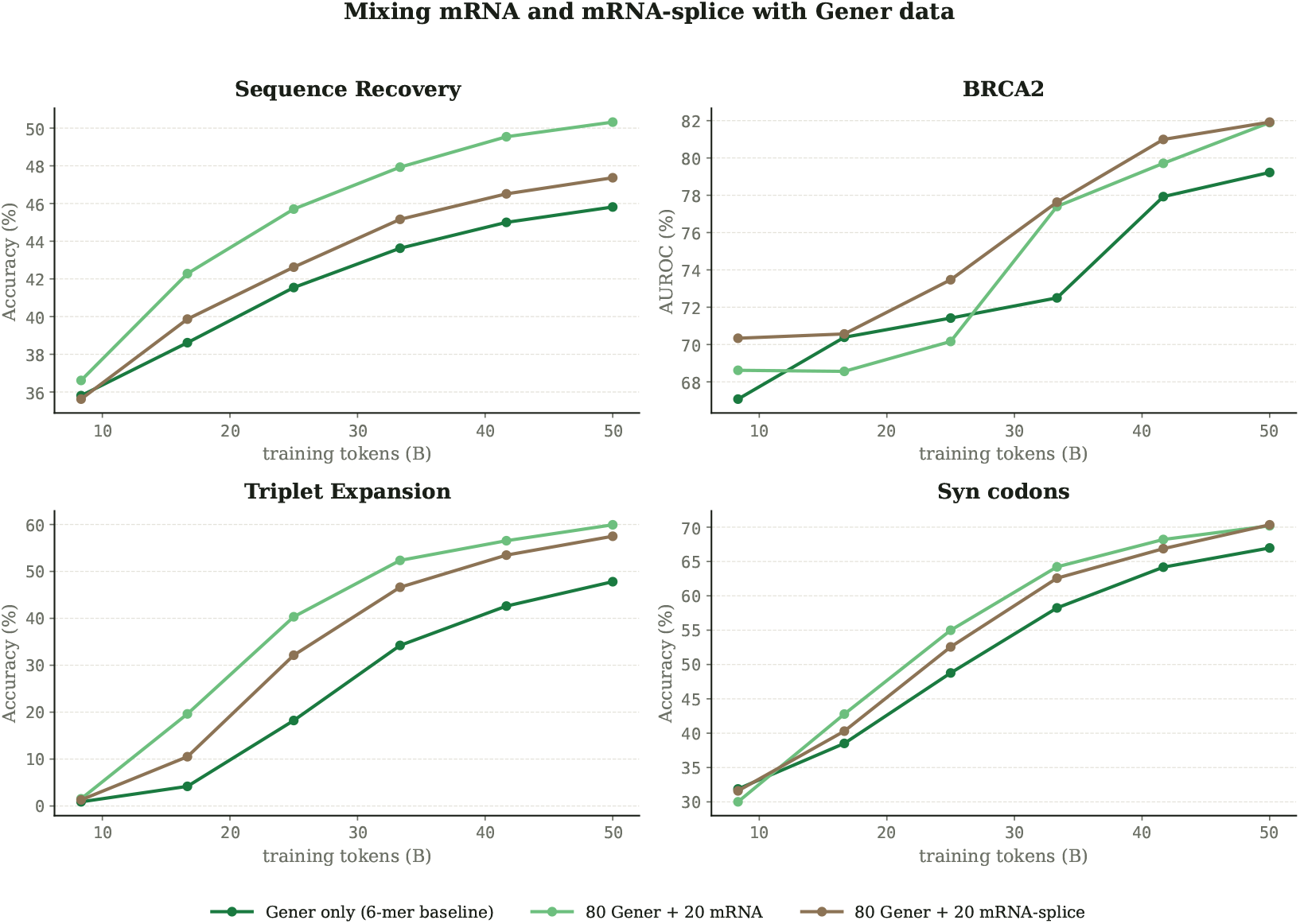
Adding 20% transcript-level data (mRNA or mRNA-splice from OpenGenome2) to the GENERator backbone. Both improve every metric over the GENERator-only baseline. mRNA gives the largest gain on SR (+4.5 pp) and triplet-expansion (+12 pp); on BRCA2 and synonymous codon the two mixes are essentially tied. The final Car-bon mixture allocates the larger share to mRNA (16%) and a smaller share to mRNA-splice (4%).

**Figure S6:**
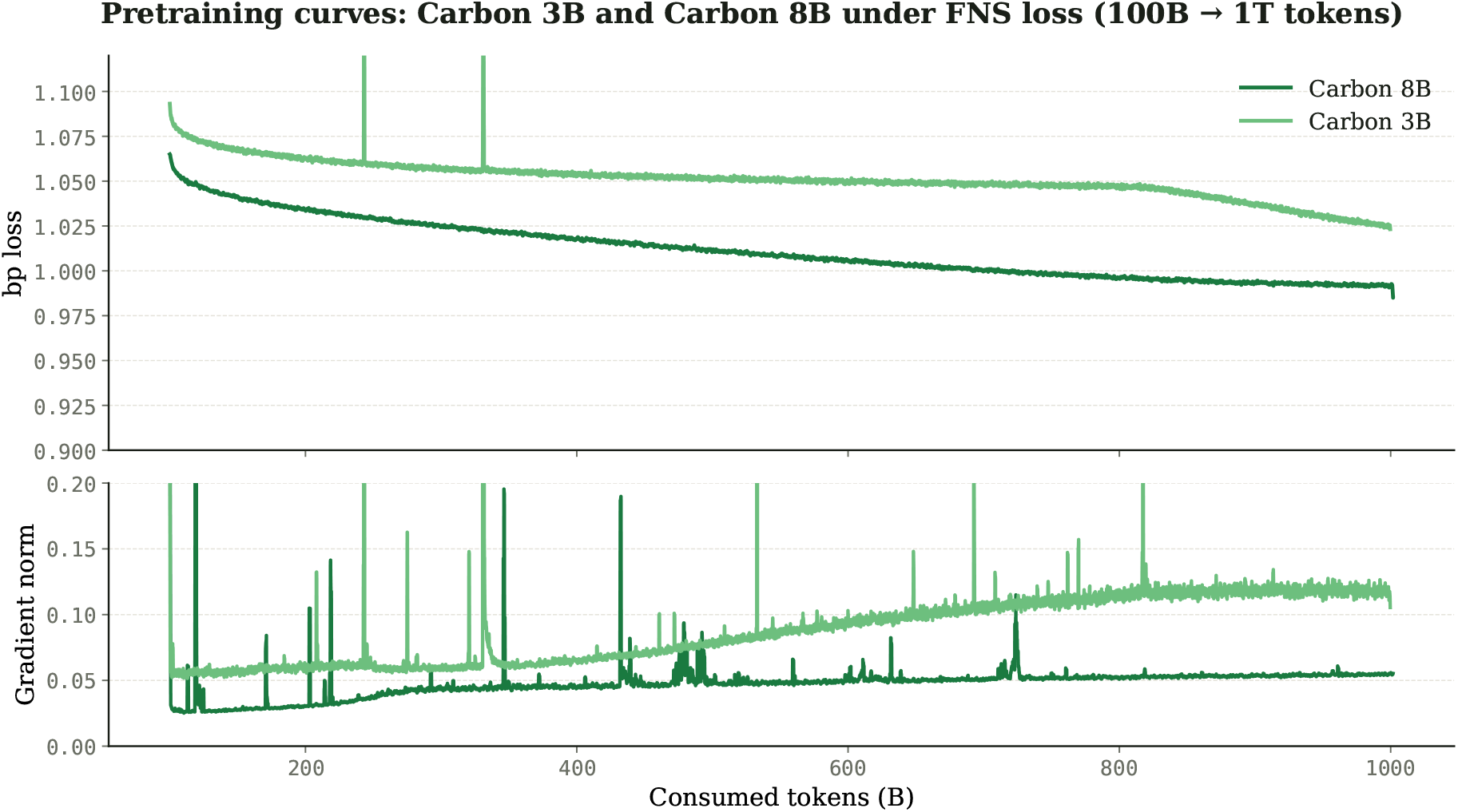
FNS-phase training loss for Carbon-3B and Carbon-8B from the CE-to-FNS switch at 100 B tokens through the end of pretraining at ~ 1 T tokens. Both models show stable training under FNS, without the loss staircases observed under prolonged CE training. The two curves take different shapes in the late-training portion by design: Carbon-3B follows a Warmup-Stable-Decay (WSD) schedule with a linear decay phase, while Carbon-8B follows a cosine decay schedule. Gradient norms remain stable throughout the FNS phase for both models, in contrast to the steady pre-staircase rise observed under CE training.

## D Finetuning

We conducted supervised fine-tuning experiments to test whether pretrained Carbon models can be adapted from next-token genomic sequence modeling to quantitative prediction of regulatory activity. The section covers three sequence-to-function regression settings: DeepSTARR enhancer activity, Random Promoter DREAM 2022 promoter activity, and Malinois/Gosai MPRA enhancer activity. For Carbon results, we report Pearson correlation coefficient (pcc; Pearson’s *r*) as the primary metric and Spearman’s *ρ* as a secondary rank-correlation metric when available. Mean pcc denotes the arithmetic mean across assay targets.

These are supervised transfer experiments rather than a single standardized benchmark. For each external result, the text states whether the comparison uses the same metric and held-out split, or whether differences in pre-processing, target transforms, checkpoint provenance, or challenge scoring make the numbers non-equivalent.

### D.1 DeepSTARR Enhancer Activity

DeepSTARR enhancer activity prediction tests how well DNA sequence predicts reporter-measured regulatory activity in Drosophila melanogaster S2 cells [67]. We fine-tuned Carbon on the Hugging Face Hub dataset repository `GenerTeam/DeepSTARR-enhancer-activity`, using 402,296 train examples, 40,570 validation examples, and 41,186 held-out test examples. The regression head was trained on the dataset-provided scaled targets `Dev_log2_enrichment_scaled` and `Hk_log2_enrichment_scaled`. Figure S7 shows observed-vs-predicted plots report log_2_ activity for the Dev and Hk targets.

Full fine-tuning with a mean-squared-error (mse) sequence regression head produced the best DeepSTARR result in our sweep. The selected Carbon-3B mse run reached test mean pcc 0.748, with Dev pcc 0.706 and Hk pcc 0.791. This is 0.038 mean pcc above the original DeepSTARR CNN reference and 0.007 below the GENERator activity predictor reported for the DeepSTARR held-out setting [67, 10].

**Figure S7:**
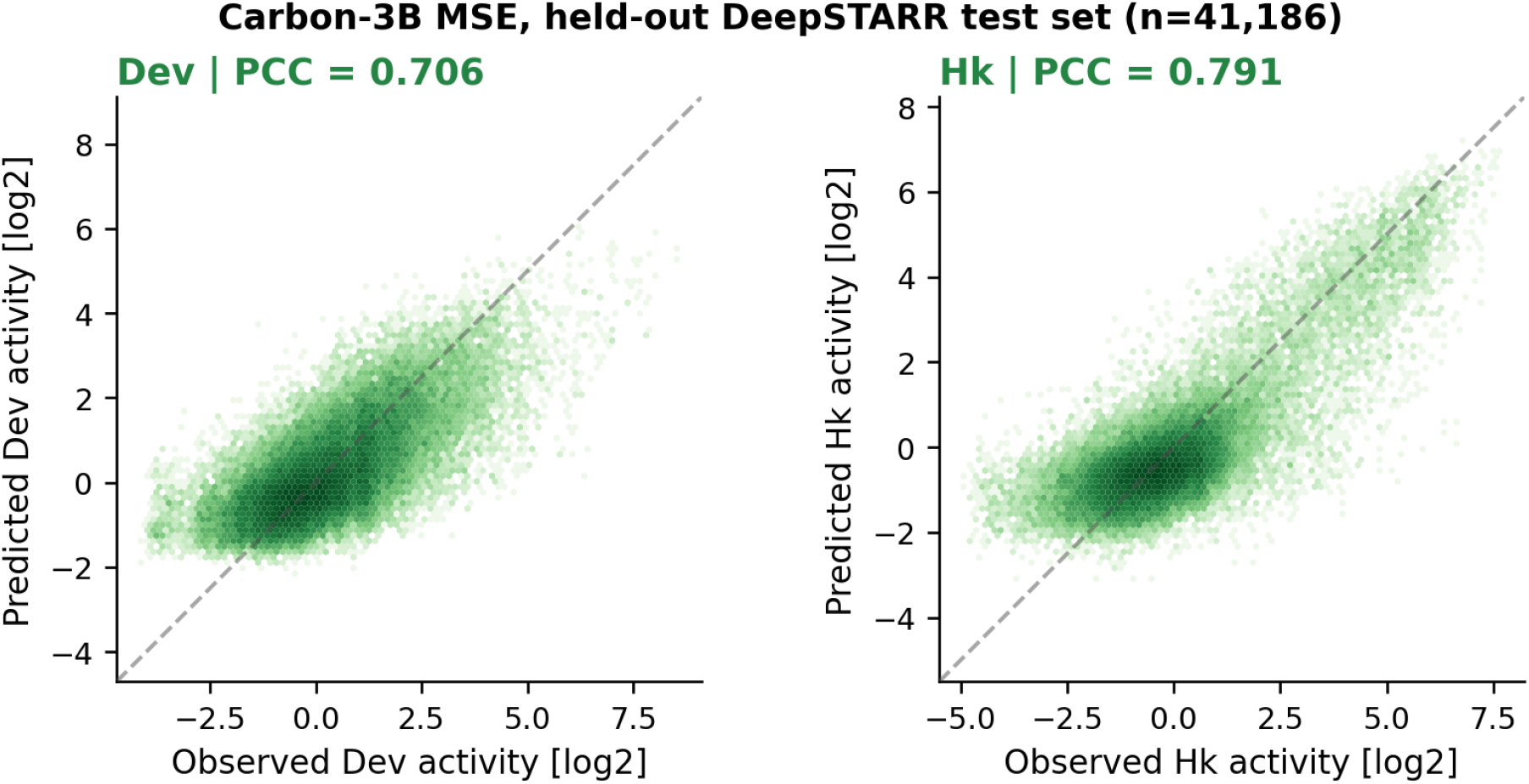
DeepSTARR observed-vs-predicted activity. Observed-vs-predicted log_2_ activity on the held-out Deep-STARR test set for the selected Carbon-3B mse model (*n* = 41,186). pcc is computed separately for Dev and Hk.

Overall, the best Carbon run is close to the strongest published DeepSTARR reference we compared against, while still trailing the task-specific GENERator activity predictor by a small margin.

### D.2 Random Promoter DREAM Activity

The Random Promoter DREAM Challenge tests quantitative expression prediction from synthetic promoter sequences [69, 70]. We converted the processed challenge data into the derived Hugging Face Hub dataset repository `HuggingFaceBio/random-promoter-dream-2022`, then fine-tuned Carbon models with a scalar regression head on a 200k-example training subset. The run reported below used DNA-mode tokenization and a correlation-oriented Pearson-Huber objective:

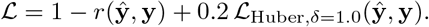

**Table S2:** DeepSTARR held-out test performance. Means are computed from full-precision run metrics and then rounded.

| Method/model | Dev PCC | Hk PCC | Mean PCC | Source |
| --- | --- | --- | --- | --- |
| ExplaiNN | 0.61 | 0.71 | 0.66 | Published baseline [68] |
| DeepSTARR | 0.68 | 0.74 | 0.71 | Original CNN benchmark [67] |
| CARBON-500M | 0.661 | 0.768 | 0.715 | MSE fine-tune |
| CARBON-3B | 0.706 | 0.791 | 0.748 | MSE fine-tune; Figure S7 |
| CARBON-8B | 0.701 | 0.790 | 0.745 | MSE fine-tune |
| GENERator-3B | 0.71 | 0.80 | 0.755 | Table S7 [10] |

Here *r*(**ŷ, y**) is Pearson’s correlation over the scalar predictions and labels in the optimization batch, and the Huber term is mean-reduced on the same train-standardized values. In distributed runs, the implementation gathers prediction and label tensors across ranks before computing the Pearson term.

The best Carbon run reached plain full-test Pearson correlation *r* = 0.807, squared Pearson correlation *r*^2^ = 0.651, and Spearman correlation *ρ* = 0.806 on the 71,103-sequence labeled test file. These values are not official DREAM leaderboard scores: the published challenge score is a weighted aggregate across subsets and includes paired-sequence subsets. The comparison to the DREAM reference model and Autosome.org is therefore not leaderboard-equivalent: our value is a plain correlation on the labeled test file, whereas the DREAM leaderboard used a weighted aggregate score across multiple subsets [69].

**Figure S8:**
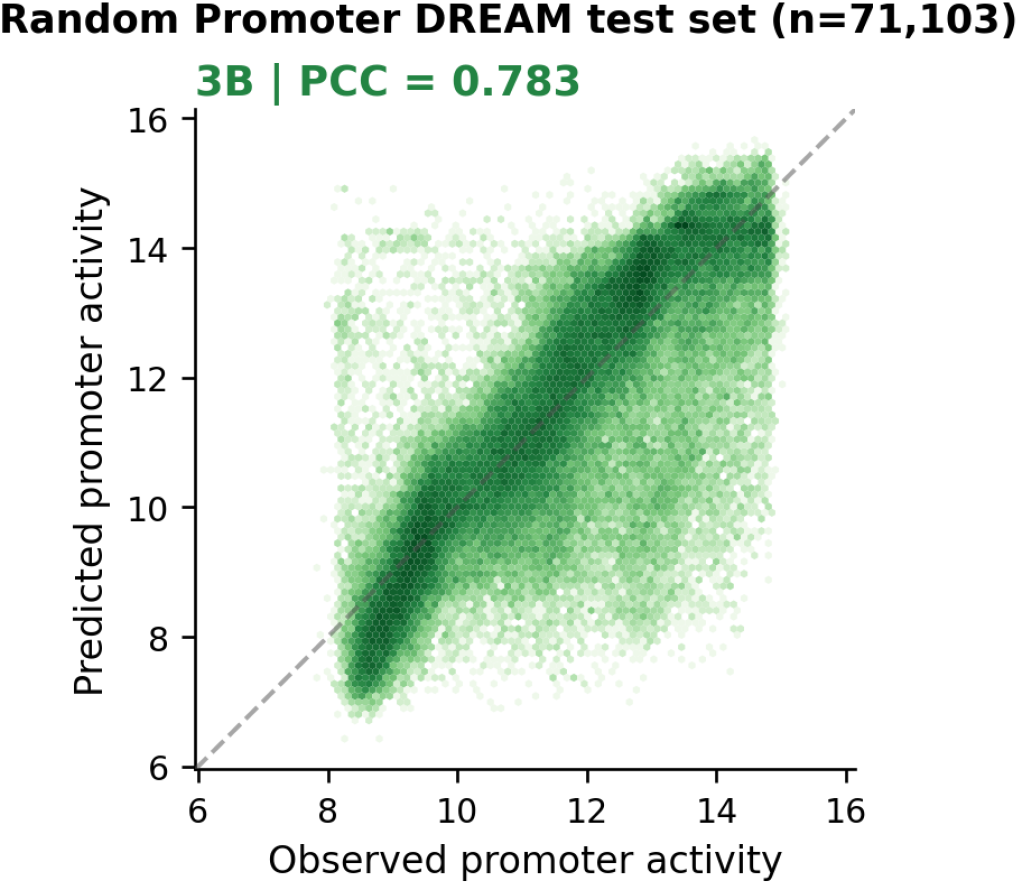
Random Promoter DREAM observed-vs-predicted activity. Observed-vs-predicted promoter activity on the held-out Random Promoter DREAM test set for the selected Carbon-3B model (*n* = 71,103).

The promoter result shows that Carbon can learn this scalar regulatory-activity signal from a modest supervised subset, but it should be interpreted as a plain-correlation transfer result rather than as a replacement for the official DREAM scoring protocol.

### D.3 Malinois MPRA Activity

The Malinois benchmark evaluates how well models predict cell-type-specific regulatory activity from synthetic DNA elements, using MPRA measurements in K562, HepG2, and SK-N-SH cells [18]. [71]. We fine-tuned Carbon on the same three log_2_ fold-change targets using the Gosai et al. MPRA supplementary data, with chromosome holdouts matching the Malinois Figure 1c setup: validation on chromosomes 19, 21, and X, and test on chromosomes 7 and 13. The selected recipe used full-model fine-tuning with a sequence regression head, mse on train-z-scored labels, DNA-mode tokenization, reverse-complement training augmentation, and reverse-complement averaging at evaluation. Figure S9 shows bbserved-vs-predicted log_2_ fold-change on the chromosome 7/13 Malinois test set.

**Table S3:**
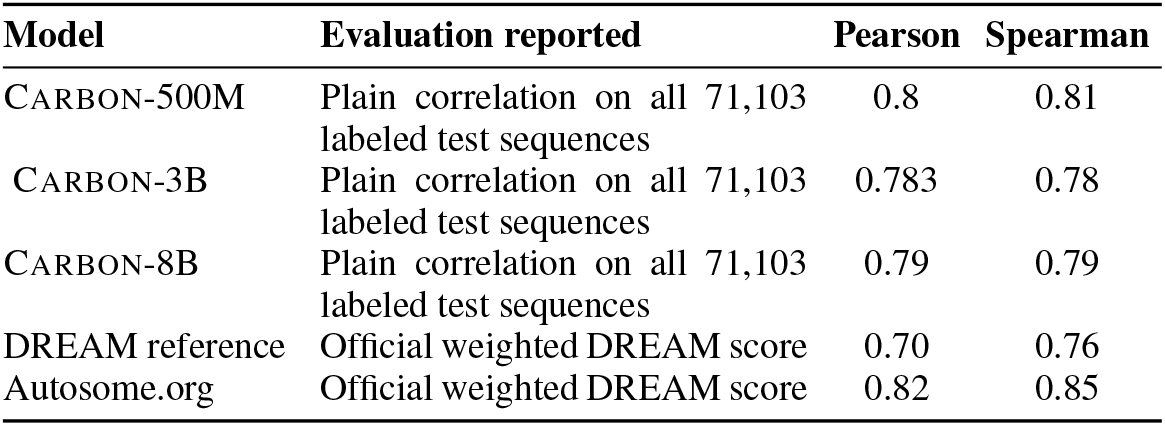
Random Promoter DREAM comparison. The Carbon rows report plain full labeled test-set correlations, whereas the challenge rows report official weighted DREAM scores.

| Model | Evaluation reported | Pearson | Spearman |
| --- | --- | --- | --- |
| CARBON-500M | Plain correlation on all 71,103 labeled test sequences | 0.8 | 0.81 |
| CARBON-3B | Plain correlation on all 71,103 labeled test sequences | 0.783 | 0.78 |
| CARBON-8B | Plain correlation on all 71,103 labeled test sequences | 0.79 | 0.79 |
| DREAM reference | Official weighted DREAM score | 0.70 | 0.76 |
| Autosome.org | Official weighted DREAM score | 0.82 | 0.85 |

**Figure S9:**
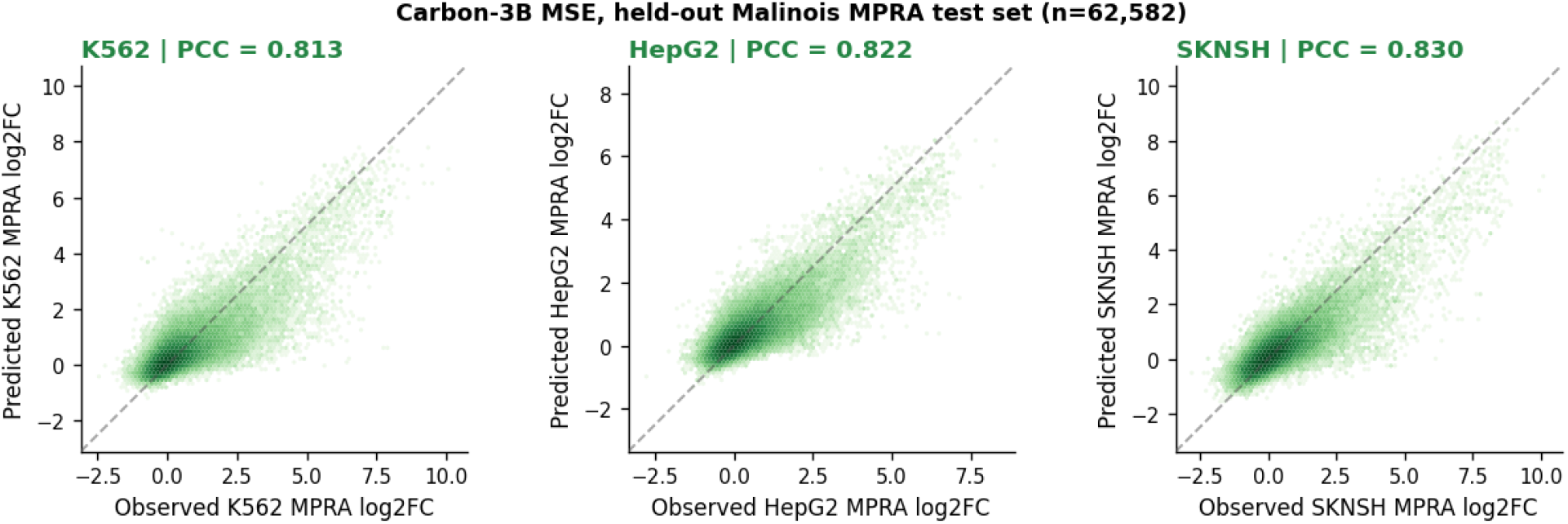
Malinois MPRA observed-vs-predicted activity. Observed-vs-predicted log_2_ fold-change on the chromosome 7/13 Malinois test set for the selected Carbon-3B mse checkpoint (*n* = 62,582). pcc is computed separately for K562, HepG2, and SK-N-SH.

Carbon-3B had the highest mean pcc on both validation and held-out test; the matched 3B test rerun reached mean pcc 0.8217 and mean Spearman *ρ* = 0.7233.

**Table S4:**
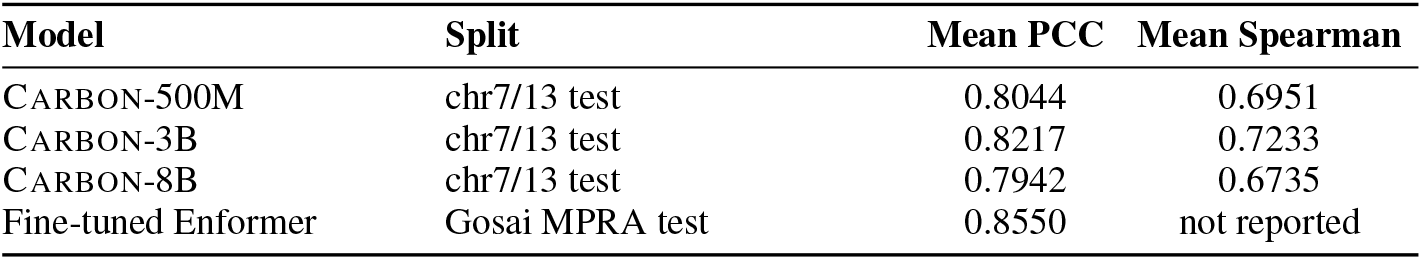
Malinois validation and held-out test comparison. The Enformer number comes from a public model card; this table should not be read as a strict rank ordering until identical preprocessing and split handling are verified.

Under this recipe, Carbon-3B had the highest Carbon held-out test mean pcc, exceeding Carbon-500M by 0.0173 and Carbon-8B by 0.0275. It remains about 0.033 mean pcc below the public fine-tuned Enformer comparison [72]; because we have not verified identical preprocessing and split handling, we do not treat the Enformer and Carbon rows as a strict rank ordering. The main positive result is that a simple full-model Carbon fine-tune recovers strong MPRA activity signal across all three cell types.

### D.4 Fine-Tuning Protocol

Across these tasks, the reported runs used full-model fine-tuning with a lightweight sequence-level regression head and validation-based checkpoint selection. All reported runs used AdamW [73], zero weight decay, 5% warmup, maximum tokenized length 512, and DNA-mode tokenization. For raw DNA inputs, `auto_dna_tags=True` automatically wraps sequences with DNA boundary tags so that the Carbon tokenizer applies DNA k-mer tokenization rather than generic BPE tokenization. Batch sizes in Table S5 are global training batch sizes.

**Table S5:** Key hyperparameters for the reported fine-tuning runs.

| Task | Hyper-parameters |
| --- | --- |
| DeepSTARR | MSE; learning rate $2 \times 10^{-5}$; batch 64; 3 epochs; reduce-on-plateau scheduler |
| Random Promoter DREAM | Pearson-Huber; learning rate $2 \times 10^{-5}$; batch 16; 1 epoch; cosine scheduler |
| Malinois/Gosai MPRA | MSE; learning rate $1 \times 10^{-5}$; batch 256; 3 epochs; reduce-on-plateau scheduler |

## Footnotes

1 The full corpus is released at https://huggingface.co/datasets/HuggingFaceBio/carbon-pretraining-corpus.

2 Evaluation code: https://github.com/huggingface/carbon/tree/main/evaluation.

3 We deliberately disable chunked prefill to measure intrinsic per-request memory cost rather than savings from engine-level optimizations. Chunked prefill is enabled for Evo2-7B only in our Genomic-NIAH evaluation (Table 5), without which it cannot reach those context lengths at all.

